# Ultrahigh Resolution Lipid Mass Spectrometry Imaging of High-Grade Serous Ovarian Cancer Mouse Models

**DOI:** 10.1101/2023.10.30.564760

**Authors:** Xin Ma, Andro Botros, Sylvia R. Yun, Eun Young Park, Olga Kim, Ruihong Chen, Murugesan Palaniappan, Martin M. Matzuk, Jaeyeon Kim, Facundo M. Fernández

**Affiliations:** School of Chemistry and Biochemistry, Georgia Institute of Technology, Atlanta, Georgia 30332, United States; Departments of Biochemistry and Molecular Biology, Indiana University School of Medicine, Indiana University Melvin and Bren Simon Comprehensive Cancer Center, Indianapolis, Indiana, 46202, United States; Department of Pathology & Immunology, Baylor College of Medicine, Houston, TX 77030, United States; Center for Drug Discovery, Department of Pathology & Immunology, Baylor College of Medicine, Houston, TX 77030, United States; Petit Institute of Bioengineering and Bioscience, Georgia Institute of Technology, Atlanta, Georgia 30332, United States

**Keywords:** Mass spectrometry imaging, matrix-assisted laser desorption/ionization, high-grade serous ovarian cancer, lipidomics, biomarkers

## Abstract

No effective screening tools for ovarian cancer (OC) exist, making it one of the deadliest cancers among women. Considering little is known about the detailed progression and metastasis mechanism of OC at a molecular level, it is crucial to gain more insights on how metabolic and signaling alterations accompany its development. Herein, we present a comprehensive study using ultra-high-resolution Fourier transform ion cyclotron resonance matrix-assisted laser desorption/ionization (MALDI) mass spectrometry imaging (MSI) to investigate the spatial distribution and alterations of lipids in ovarian tissues collected from double knockout (*n* = 4) and a triple mutant mouse models (*n* = 4) of high-grade serous ovarian cancer (HGSC). Lipids belonging to a total of 15 different classes were annotated and their abundance changes compared to those in healthy mouse reproductive tissue (*n* = 4), mapping onto major lipid pathways involved in OC progression. From intermediate-stage OC to advanced HGSC, we provide a direct visualization of lipid distributions and their biological links to inflammatory response, cellular stress, cell proliferation, and other processes. We also show the ability to distinguish tumors at different stages from healthy tissues *via* a number of highly specific lipid biomarkers, providing targets for future panels that could be useful in diagnosis.

## Introduction

Ovarian cancer (OC) is one of the most lethal cancers among women, with patients suffering from the highest mortality rate among all gynecological cancer patients.(Kandimalla et al., 2021, Cabasag et al., 2022) Due to the lack of symptoms at its early (localized) stages, only a small portion of cases is diagnosed early enough for effective treatment.(Dilley et al., 2020) Current diagnostic tools, including transvaginal ultrasound and cancer antigen (CA)-125 blood tests do not provide sufficient sensitivity and specificity, especially for early stage OC diagnosis.(Kamal et al., 2018) Therefore, neither of the aforementioned methods are used as screening tools. Among all OC subtypes, high-grade serous ovarian cancer, also known as high-grade serous carcinoma (HGSC), causes 70 – 80% of all OC deaths,(Lisio, 2019) as it is typically diagnosed at distant (late) stages. A more in-depth understanding of the molecular pathogenesis of HGSC could help save lives by providing targets of diagnostics and prognostic value.

Lipids play crucial roles in cancer pathogenesis and are critical effectors in energy storage, cell signaling, and maintaining cell structures.(Butler et al., 2020, Ma and Fernández, 2022) Abnormal alterations in lipid levels in the tumor microenvironment usually follow cancer progression, making them informative cancer markers.(Butler et al., 2020) Lipid alterations in biofluids such as serum and plasma due to HGSC pathogenesis have been measured *via* liquid chromatography–tandem mass spectrometry (LC-MS/MS), in both animals and humans.(Sah, 2022, Li et al., 2016, Cheng et al., 2020, Mir et al., 2021) These studies provide detailed descriptions of how alterations of various lipid classes are reflected in various compartments during OC development, but fail to capture the specific spatial lipid distributions as they are present in tissue. Spatially resolved lipidomics can provide such maps in tissues and organs, showcasing the heterogeneity of cancer tumors and any uncommon lipids that could potentially be shed into adjacent biofluids. These molecularly-specific maps are helpful in establishing the relationship between lipidome changes at the tissue level with lipid alterations in the surrounding biofluids.(Petras et al., 2017)

Mass spectrometry imaging (MSI) is a powerful tool to study altered metabolism in the context of cancer biology.(Petras et al., 2017, Taylor et al., 2021, Ma and Fernández, 2022) Amongst all MSI techniques, matrix-assisted laser desorption/ionization (MALDI) MSI is by far the most mature and widely used. With proper MALDI matrix selection, it provides excellent lipid class coverage with a spatial resolution as low as 1 µm. Herein, we present the first ultrahigh resolution Fourier transform ion cyclotron resonance (FTICR) MALDI MSI study on tissue sections collected from the reproductive systems of two types of HGSC mouse models, triple mutant (TKO) *p53*^LSL-R172H/+^ *Dicer1*^flox/flox^ *Pten*^flox/flox^ *Amhr2*^cre/+^ (Kim et al., 2015) and double knockout (DKO) *Dicer1*^flox/flox^ *Pten*^flox/flox^ *Amhr2*^cre/+^ mice.(Kim et al., 2012) These murine models reproduce human HGSC with high fidelity and therefore have translational value.(Kim et al., 2020) The ultrahigh mass resolution and excellent mass accuracy yielded by FTICR MSI enabled the creation of lipid ion maps with exquisite specificity. Lipid distributions were compared between cancerous and healthy tissues for lipids putatively annotated with low false discovery rates (FDR < 10%). The findings reported here lead to a more comprehensive understanding of lipidome remodeling associated with HGSC and produce potential HGSC lipid biomarkers that could be translated to human studies.

## Materials and Methods

### Chemicals

MALDI matrix 1,5-diaminonaphthalene (1,5-DAN, ≥ 97%), tissue embedding media gelatin from bovine skin (type B), sodium carboxymethyl cellulose (CMC), and isopentane (≥ 95%), acetonitrile, ethanol, methanol, and water for H&E staining were purchased from Sigma Aldrich (St Louis, MO). Histological-grade xylenes were purchased from Spectrum Chemical. Hematoxylin and eosin were purchased from Cancer Diagnostics, Inc. and FisherBrand (Pittsburgh, PA), respectively. All chemicals were used as received.

### Animal generation, tissue collection, preservation, and sectioning

The animals used in this study included double-knockout (DKO) mice and triple mutant (TKO) mice (collected at an advanced stage). Matched controls were generated following protocols previously described.(Paine et al., 2016, Sah, 2022) All animals were sacrificed in accordance with animal protocol #21124 approved by the IACUC at Indiana University School of Medicine (Indianapolis, IN, USA). Whole reproductive systems collected from TKO, DKO and control mice were embedded in a 1% CMC and 5% gelatin aqueous solution. The embedding temperature was maintained at −20 °C using an isopentane-dry ice bath. Embedded tissues were stored at −80 °C until further use and sectioned using a CryoStar NX70 Cryostat operated at −20 °C. The thickness of the sectioned tissue slices was kept at 10 μm. Slices were transferred to Fisherbrand™ Superfrost™ Plus microscope slides immediately and stored for MALDI MSI experiments.

### Matrix deposition

1,5-DAN was dissolved in 90/10 ACN/water (v/v) to yield 5 mg mL^-1^ solution. Tissue sections were sprayed with the MALDI matrix solution by using an HTX TM-Sprayer^TM^ (Chapel Hill, NC). This sprayer was operated using the following parameters: the nozzle temperature was set to 30 °C, the flow rate of the matrix solution and the N_2_ drying gas were 0.1 mL min^-1^ and 2 L min^-1^, respectively; the nozzle moved at a velocity of 1200 mm min^-1^ and the spray tracking space was 2.5 mm. The solution was sprayed in a crisscross pattern for 6 cycles. No drying time was set between each cycle.

### Mass spectrometry

A Bruker SolariX 12-Tesla FTICR mass spectrometer (Bruker Daltonics, Bremen, Germany) equipped with a MALDI ion source was used for all MSI experiments. The mass spectrometer was operated in negative ion mode in the 150 – 1200 *m/z* range. The time domain data set size for the MSI experiments was set to 4,000,000, which was equivalent to a mass resolution of 410,000 at *m/z* 400. The free induction decay (FID) transient time was 1.677 s. The MALDI laser power was set to 30%, the number of laser shots accumulated on each pixel was 300. The laser repetition frequency was 1000 Hz and the laser beam focus was set to small. The spatial resolution (defined as the scanned pixel size) for MSI experiments was set to 50 μm × 50 μm, which led to 40 – 50 hours of data acquisition time and 800 GB to 1 TB file size per tissue slice. The mass spectrometer was calibrated externally with a sodium trifluoroacetate aqueous solution and internally with the FA(18:1) and PI(38:4) lipids to ensure a mass accuracy better than 1 ppm, on average. For each animal model, 4 reproductive systems from different mice were sectioned and imaged. A minimum of 4 technical replicates were performed for each distinct reproductive system in random order. Ionic signals were normalized to the total ion current within each image. The mass isolation window used for selecting extracted ion images was ± 0.001 Da.

### H&E staining

Following MSI experiments, the matrix was washed off by 100% ethanol for hematoxylin and eosin (H&E) staining. For staining purposes, slides were immersed in the following solvents/solutions in a sequential fashion: 95% ethanol for 30 s, 70% ethanol for 30 s, water for 30 s, hematoxylin stain for 2 min, water for 30 s (repeat twice in different jars), 70% ethanol for 30 s, 95% ethanol for 30 s, eosin for 30 s, 95% ethanol for 30 s, 100% ethanol for 1 min (×2 in different jars) and xylenes for 1 min. The stained slides were air-dried, covered with Cytoseal and coverslips for optical imaging. Optical images were obtained on a Hamamatsu NanoZoomer Scanner and exported with the Hamamatsu NDP software.

### Data processing

The measured accurate masses for features in the mean mass spectra were subject to Lipid Maps and HMDB database searches for putative lipid annotations using METASPACE.(Palmer et al., 2017) Annotated features with an FDR < 10% were chosen for further analysis. This list of annotated features was then used for spatial segmentation, which was performed using a bisecting k-means algorithm (Alexandrov, 2012) with strong image denoising, using the SCiLS Lab software (version 2023a Pro, Bruker Daltonics, Bremen, Germany). Segmentation results were compared to H&E-stained microscopic images and used iteratively to select regions of interest (ROI). Tumor regions in TKO and DKO tissue sections and healthy ovaries and fallopian tubes were selected as ROI for further analysis. Lipid feature lists were exported for each ROI and used for the development of statistical models and for lipid pathway analysis. Specifically, the average lipid peak areas in each ROI were exported into MetaboAnalyst to perform univariate receiver operating characteristic (ROC) analysis to select potential cancer biomarkers based on their area under the curve (AUC).(Fawcett, 2006) Features with AUC ≥ 0.80 were chosen and used to build principal component analysis (PCA) models for discriminating HGSC from controls. Peak areas were normalized to the total ion current and auto scaled prior to statistical analysis. Log_2_-fold changes for all annotated features were calculated and plotted prior to lipid pathway analysis. A pathway map was generated using lipid pathway enrichment analysis (LIPEA)(Acevedo et al., 2018) together with literature reports, which provided a rationale for the observed alterations in different lipid classes and interactions between different lipids.

## Results and Discussion

### Lipid feature annotations and ROI selection

A total of 228 lipid features belonging to 15 lipid classes were putatively annotated with an FDR <10%. Approximately one third of the annotated lipids belonged to the phosphatidylethanolamine and ether-linked phosphatidylethanolamine classes, in the 450 – 800 *m/z* range. Other significant annotated lipid classes included sphingolipids (sphingomyelins, ceramides and ceramide phosphates) in the 550 – 800 *m/z* range, fatty acids in the 250 – 370 *m/z* range, phosphatidic acids in the 400 – 750 *m/z* range, phosphatidylserines in the 750 – 900 *m/z* range and phosphatidylglycerols and phosphatidylinositols in the 750 – 950 *m/z* range. For each tissue section studied, regions of interest were selected with the assistance of a spatial segmentation algorithm and co-registration with hematoxylin & eosin (H&E)-stained optical images. For TKO tissue sections, advanced-stage high-grade serous carcinoma (HGSC) had developed in all 4 biological replicates studied; ROI corresponding to these HGSC regions were selected and confirmed by spatial segmentation and H&E staining (Figure 1). Region #3 of TKO-2 (Figure 1) presented not only a HGSC region, but also a blood-filled cyst. This part of the tissue was excluded from the HGSC ROI as its molecular composition was very different from the HGSC regions, as indicated by segmentation analysis. TKO mice present the most aggressive form of HGSC, due to the *p53* mutation that results in a mean survival of 6.6 months.(Kim et al., 2020) All studied TKO tissues were collected at the 85% – 100% lifetime (defined as the ratio of the mouse age when sacrificed and the mean TKO mice survival lifetime, see Table S1 for detailed mice collection information). The lipid phenotype of HGSC tumors in TKO animals was viewed as the most advanced and aggressive compared to DKO and control animals.

**Figure 1.**
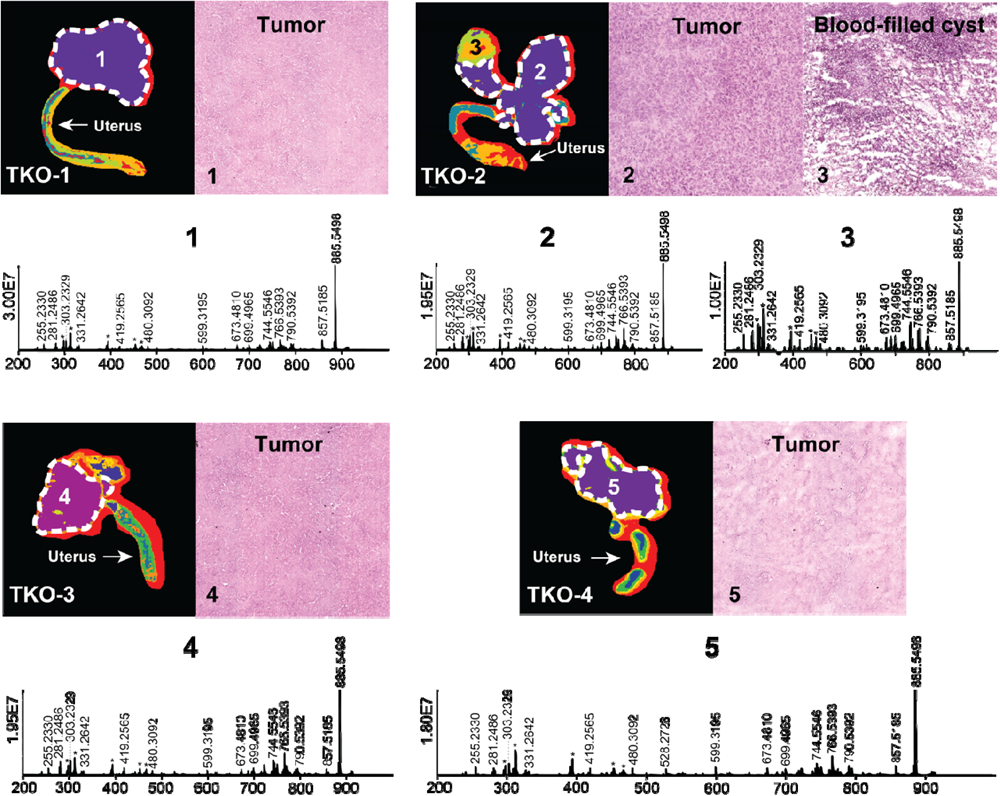
Spatial segmentation and selected ROI for reproductive tissue sections of four different TKO mice. ROI are outlined with white dashed lines. Optical images of H&E-stained selected sub-regions (labelled with numbers) are shown for each tissue sample, The mean ROI mass spectrum of each of the labelled region are also shown. Background ions are labeled with asterisks.

HGSC in DKO mice is less aggressive compared to TKO mice due to the preservation of the *p53* gene.(Kim et al., 2015) Previous studies have shown that TKO mice die earlier than DKO mice and the median survival for DKO mice is 9.1 months, 2.5 months longer than that of TKO mice.(Kim et al., 2020) DKO tissues used in this study were sacrificed at two different time points, two of them (DKO-1 and DKO-2 in Figure 2) sacrificed 10 days earlier than the other two (DKO-3 and DKO-4 in Figure 2), with a %lifetime of 77% and 83%, respectively. Therefore, stages of OC in these DKO tissues were significantly different from each other and from TKO tissues.(Table S1). *p53* mutations have been found in 96% of HGSC in human OC cases, leading to enhanced tumor aggressiveness and probability of metastasis, as observed in TKO animals.(Rivlin et al., 2011, Brosh and Rotter, 2009) Tumors in DKO tissues were essentially less aggressive due to the preserved normal *p53* functions. Based on tumor aggressiveness, histology of the DKO tissue sections and the %lifetime of the mice at the time of sacrifice, two sub-groups were identified: DKO-I (DKO-1 and DKO-2 animals where intact ovaries were retained and yet to be invaded by the HGSC tumor, Regions 7 and 10 in DKO-1 and DKO-2, respectively) and DKO-II (advanced-stage tumors including the DKO-3 and DKO-4 animals). H&E optical images of DKO-I tissue sections revealed that several blood-or fluid-filled cysts had developed adjacent to the tumor region (Region 6, a thin region around the cyst where obvious cell proliferation was observed in the H&E image), resulting in a highly heterogeneous morphology compared to DKO-II and TKO tissues. Cysts were excluded from HGSC ROIs and their lipidomic profiling is discussed separately. In fact, we combined intact ovaries as part of the control tissues (healthy fallopian tubes and ovaries) and developed multivariate statistical models for differentiation of HGSC from healthy tissues. Figure 2 shows segmented MS images, H&E images and mean mass spectra for DKO-I and DKO-II tissues. Blood-filled cysts (*e.g.* region 7 in Figure 2) showed significant differences in the abundances of measured lipid ions compared to the adjacent tumor regions (Region 6 in DKO-1), especially the lipid ions in the 680 – 820 *m/z* range, mainly consisting of phosphatidylethanolamine (PE), phosphatidylserine (PS) and phosphatidylinositol (PI).

**Figure 2.**
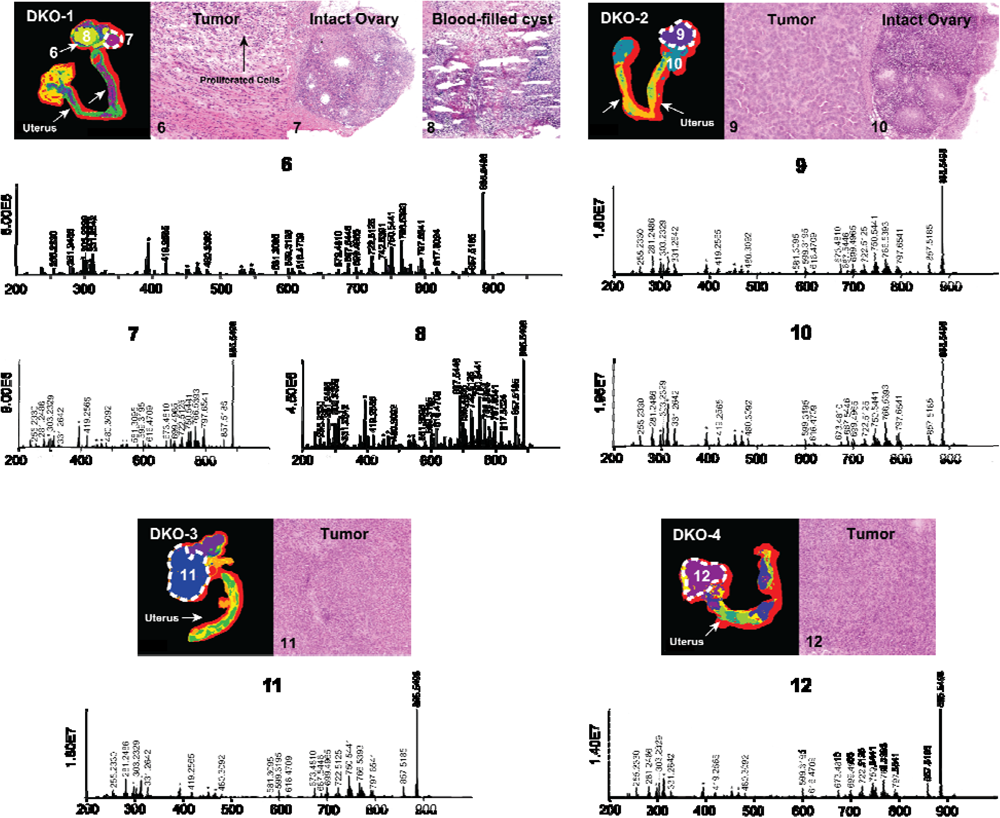
Spatial segmentation and selected ROI for reproductive tissue sections of four different DKO animals. The ROI are outlined with white dashed lines. Optical images of H&E-stained selected sub-regions (labelled with numbers) are shown for each tissue sample, the mean ROI mass spectra of each of the labelled region are also shown. Background ions are labeled with asterisks.

### Multivariate analysis

To examine the potential of the annotated lipid features as discriminant OC tissue markers, we conducted univariate receiver operating characteristic (ROC) analysis to select the most promising lipids from the 228 annotated features. ROC curves (sensitivity vs. 1 – specificity) were plotted and the area under the curve (AUC) was calculated for each annotated lipid. Lipid features with AUC ≥ 0.80 were considered acceptable and preliminary selected.(Mandrekar, 2010) For differentiation of DKO *vs*. control tissues (healthy fallopian tubes and ovaries), 92/228 lipid features passed this filter. For TKO vs. control tissues, 108 lipid features were picked. 152 and 177 lipid features were selected by univariate ROC while DKO and TKO tumors were compared to cysts in DKO tissues. Table S2 shows the features’ AUC values, *p*-values and log_2_-fold changes (FC). In addition to compare HGSC to healthy fallopian tubes and ovaries, we profiled the significantly altered lipids in cysts in the DKO-1 tissues and compared lipid alterations to HGSC regions. Filtered lipid feature lists were used to develop principal component analysis (PCA) models to examine the capabilities of these lipid panels for differentiating tumor regions from healthy tissues (Figure 3 (a) and 3 (b)) or non-tumor regions (necrotic cysts) in cancerous tissues (Figure 3 (c) and 3 (d)). As illustrated in Figure 3, each tumor sub-group led to highly distinct clusters and was successfully separated from the cluster of healthy controls or necrotic cysts. For TKO tumors, tight clusters formed, suggesting the high homogeneity of the advanced-stage HGSC. For DKO clusters, some sub-cluster finer structures were observed within the groups, revealing differences between biological replicates (labeled in Figure 3 (a) and 3 (c)).

**Figure 3.**
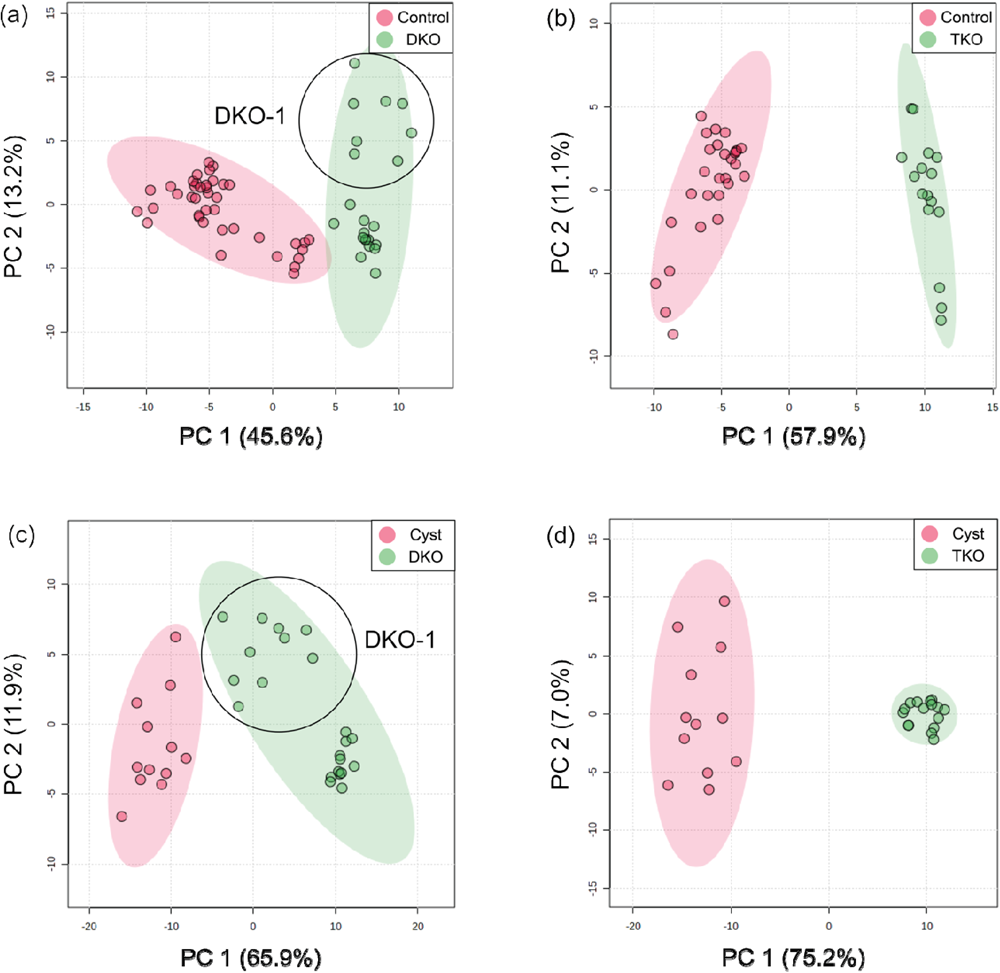
PCA score plots for differentiation of (a) DKO and (b) TKO tumors from ovaries and fallopian tubes in healthy tissues (controls), and differentiation of (c) DKO and (d) TKO tumor from cysts (non-tumor regions in tumor tissues). All PCA plots were constructed using the lipid features filtered by their AUC values by ROC analysis.

### Lipid alterations and their correlations with OC progression

As a first approach to understand lipid pathways altered in HGSC, we investigated the alterations of all 228 annotated lipid features in tumor and control tissues by calculating log_2_FC for different lipid classes. The peak areas of the lipids within each class were summed, the log_2_FC were calculated, plotted and categorized based on tumor stage, as shown in Figure 4. Clear trends were observed for most lipid classes except for FA and ether PE. We estimated that for FA, the measured abundances may be biased due to residual fat on the tissue sections, skewing the observed relationship between FA alterations and the OC progression.

**Figure 4.**
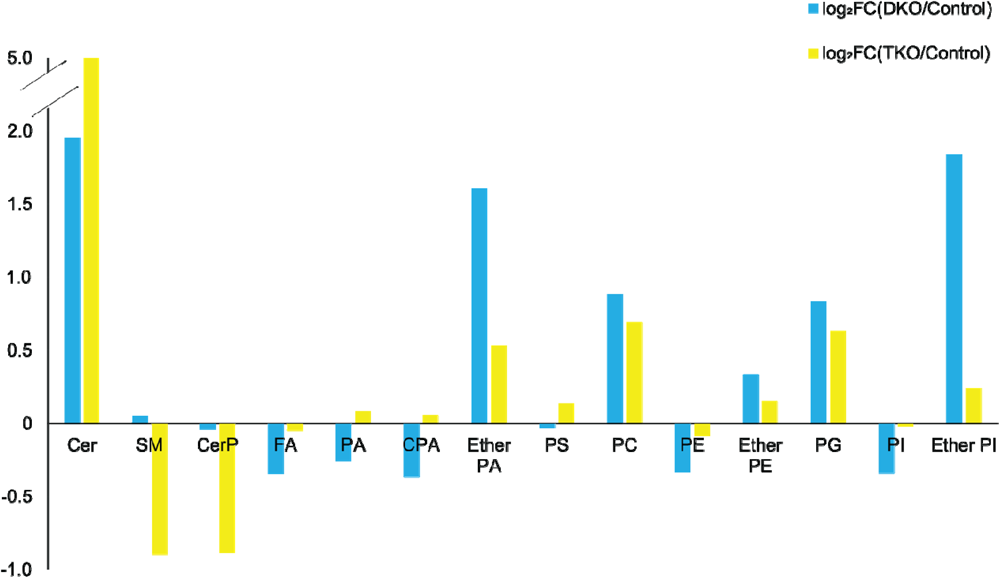
Log_2_-fold change (log_2_FC) plot for the total lipid abundances grouped by lipid classes for DKO vs. control (blue bars) and TKO vs. control (yellow bars).

A schematic lipid pathway map involving the major lipid classes detected in MSI experiments is given in Figure 5 (corresponding images and mass errors are provided in Figure 6 and S1). MS images of selected lipid ions from each lipid class are displayed in Figure 6. We discuss each lipid class depicted in Figure 4 in detail below.

**Figure 5.**
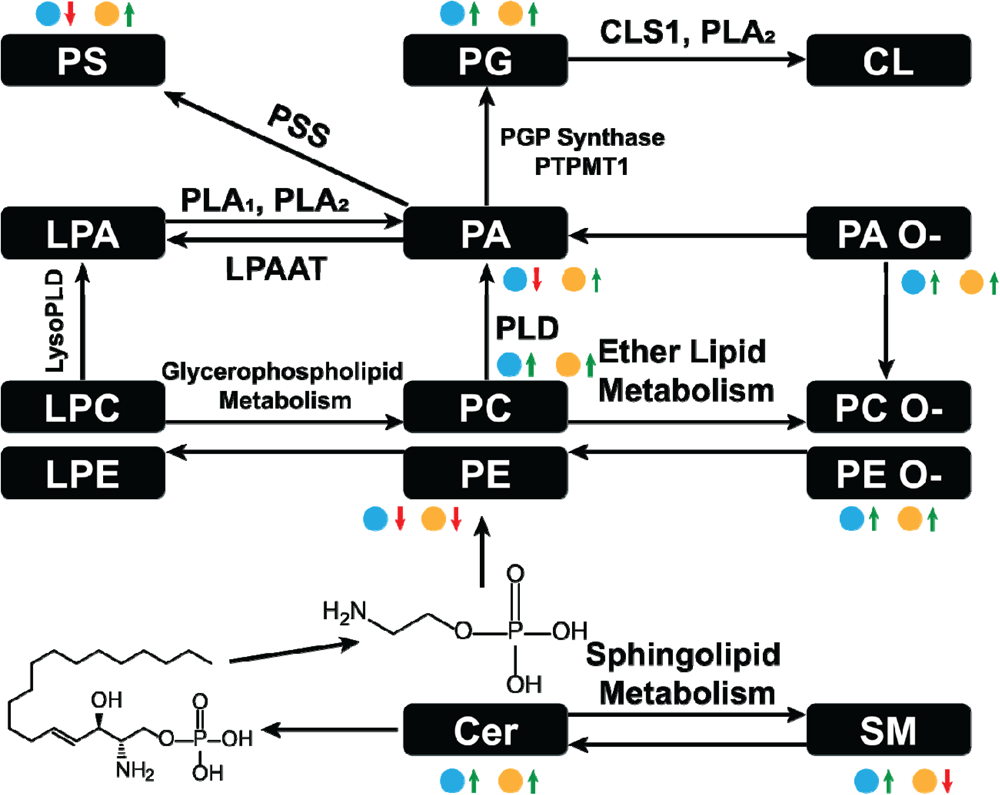
Major lipid pathways involved in OC progression as revealed by MSI experiments. Log_2_FC magnitudes are shown next to each lipid class. The blue dot represent log_2_FC(DKO/Control) and the yellow dot represents log_2_FC(TKO/Control). Red arrows indicate a negative log_2_FC and green arrows a positive log_2_FC. The more arrows, the larger the log_2_FC. PLA: phospholipase A, PLD: phospholipase, DLPAAT: lysophosphatidyl acyltransferase, PSS: L-serine phosphatidyltransferase, PGP synthase: glycerol phosphate synthase, PTPMT1: protein tyrosine phosphatase mitochondrion 1, CLS1: cardiolipin synthase 1.

**Figure 6.**
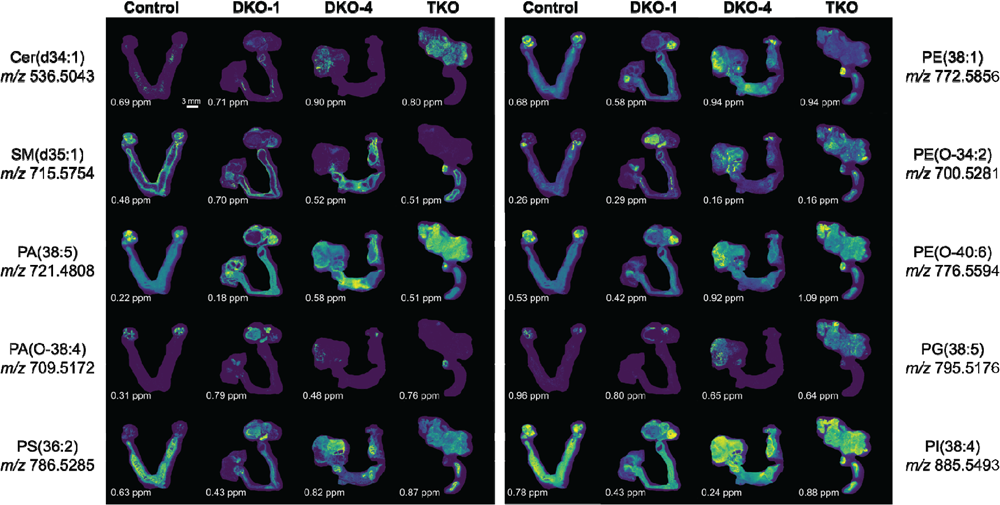
Selected extracted ion images for key lipid ions in each lipid class altered in OC. The mass errors of each detected lipid ion are displayed at the bottom left corner of each image, highlighting the high mass accuracy of the FTICR mass spectrometer employed. The scale bar of the images is provided at the bottom right corner of the control image at *m/z* 536.5043.

#### Sphingolipids

Sphingolipids, including ceramides (Cer), sphingomyelins (SM) and ceramide 1-phosphates (CerP) were drastically altered in all tumor regions, in agreement with their known roles in cell growth, survival and death during cancer progression.(Hannun and Obeid, 2008) As HGSC develops, Cer were significantly accumulated in both DKO and TKO tissues as OC progressed to HGSC (log_2_FC(DKO/Control) = 1.95 and log_2_FC(TKO/Control) = 4.78). SM showed a slight increase in DKO tissues whereas major downregulation was observed in TKO tissues. CerP were observed to decrease in both DKO and TKO tissues, indicating their conversion to Cer as HGSC develops. Cer are key substrates for many important enzymes such as ceramide synthase and ceramidase (Kreitzburg et al., 2018) and can be generated *de novo* during tumor necrosis processes occurring in cancer development.(Takabe et al., 2008) In tumor regions of DKO and TKO tissues, conversion from SM and CerP to Cer suggested the pro-mitogenic properties of OC and the increased drug resistance and proliferation probability of the cancer cells.(Sah, 2022) In particular, SM and CerP were found to be much lower in abundance in HGSC regions of TKO tissues, likely due to their conversion to Cer as a response to oxidative stress and other cellular stresses that mediate cell death.(Fekry et al., 2018) Sphingolipids alterations in advanced-stage OC tissue sections were also in agreement with previous LC-MS longitudinal studies in serum from the same animal model.(Sah, 2022) Interestingly, in DKO-1 mice, Cer were also found to accumulate in blood-filled cysts instead of tumor regions, and as they eventually transformed to HGSC, Cer started to show major accumulations (Figure 6).

#### Phosphatidic acids

Phosphatidic acids (PA) and cyclic PA (CPA) were found to decrease in DKO tissues and increase in TKO tissues compared to healthy fallopian tubes and ovaries. PA may also be generated from MALDI in-source decay of phosphatidylserines (PS).(Hu et al., 2022) Therefore, we first tested our experimental conditions by varying the laser power applied in MALDI experiments (Figure S2). A stable PS/PA ratio indicated that in source artifacts were not significant under our chosen experimental conditions. A 30% laser power was selected for all our experiments to ensure high ion abundances and signal to noise ratios with minimal PS fragmentation. As noted in Figure 5, PA serve as central precursors for biosynthesis of a variety of glycerophospholipids and are known to be a major lipid class for regulating cell proliferation.(Dória et al., 2016) Interestingly, PA were not observed in blood-filled cysts in DKO-1 tissues (Figure 6). On the contrary, ether-linked PA (PA O-) were mainly accumulated in these cysts, together with PA O-in healthy control tissues (Figure 6), indicating the conversion from PA O-to PA as OC progresses.

#### Phosphatidylserines

Phosphatidylserines (PS) are key membrane lipids involved in processes such as maintaining mitochondrial membrane integrity and neurotransmitter release.(Kaynak et al., 2022) During cancer development, PS act as signaling molecules that indicate the presence of apoptotic cells in cancer tissues.(Birge et al., 2016) PS are also responsible for immunosuppression of the tumor microenvironment, increasing the activity of dendritic cells.(Calianese and Birge, 2020) It has been observed that PS are transported from the inner cell membranes to the outer cell membranes as the cancer progresses,(Nagata et al., 2020) which agrees with our findings that PS were more abundant in the more advanced TKO tissues.

#### Phosphatidylethanolamines

Alterations in phosphatidylethanolamine (PE) metabolism are correlated with that of PS. As with PS, PE have also been found to shuttle from the inner to the outer layers of cell membranes when apoptotic and tumor cells are present.(Stafford and Thorpe, 2011) PE alterations in tumor cell membranes are known to modulate membrane protein activity, leading to dysregulated response to extracellular signals.(Kitajka et al., 2002) However, unlike previous reports of PE and PS being upregulated on cell surfaces,(Leite et al., 2015) an opposite trend was observed for these two lipid classes, i.e. PE were downregulated in tumor regions of DKO and TKO whereas PS were slightly downregulated in DKO tissues (log_2_FC = −0.02) and upregulated in TKO tissues (Figure 4), suggesting the inter-conversion between PE and PS at different OC stages. On the other hand, ether-linked PE (PE O-) were found increased (log_2_FC > 0, see Figure 4) in all tumor regions, indicating the conversion of PE to PE O-via ether lipid metabolism, which would also explain the observed PE decreases. Interestingly, PE (O-40:X) such as PE (O-40:6) (see Figure 6) showed different spatial distributions than other PE O-, and mainly accumulated in the healthy fallopian tubes and ovaries and the HGSC regions in DKO and TKO tissues. Other PE O-such as PE(O-34:2) were found localized in cysts and, comparatively speaking, much less in tumor regions.

#### Phosphatidylglycerols

We observed an accumulation of phosphatidylglycerols (PG) in tumor regions as OC develops from intermediate to advanced stages. As key intermediates of cardiolipin (CL) synthesis,(Butler et al., 2020) the upregulation of PG may indicate a significant upregulation of CL in tumor cells. Alterations in CL abundances are known to correlate to the regulation of the cell apoptosis rates.(Thorne et al., 2021) Literature reports indicate that CL formation is inversely related to that of PE in the mitochondrial inner membrane,(Böttinger et al., 2012) which may explain the opposite trends observed for PE and PG in our results. Because CL are not easily detected under the MALDI conditions used in this experiment, no definitive conclusions can be drawn.

#### Phosphatidylinositols

Overall lower levels of phosphatidylinositols (PI) were observed in all tumor stages, with the intermediate stage being the lowest and the more aggressive TKO tumors showing the highest abundances. This indicates the consumption of PI in the more aggressive types of HGSC. PI are involved in the PI3K/AKT pathway as key precursors for phosphatidylinositol phosphates (PIP).(Osaki et al., 2004) This pathway is altered significantly in various cancers including OC.(Osaki et al., 2004, Engelman, 2009) Dysregulation of this pathway facilitates cancer progression and drug resistance.(Mayer and Arteaga, 2016) Therefore, cancer treatments targeting this pathway have been the focus of much research effort, with several drugs now in clinical trials.(Mayer and Arteaga, 2016)

## Conclusions

In this study, we used ultra-high resolution mass spectrometry imaging to map the spatial distributions of lipids in ovarian cancer tissues originating from two different faithful mouse models of HGSC. A total of 228 lipids were annotated based on accurate *m/z* measurements using an FTICR mass spectrometer. Significantly altered lipids included sphingolipids that mainly reflected the response to cellular stresses induced by OC progression and (ether) phosphatidic acids with spatial distributions highly localized to tumor (PA) or the cyst regions (PA O-) linked to high rates of cell proliferation. Several other glycerol phospholipids that included phosphatidylserines, phosphatidylethanolamines, and phosphatidylglycerols were strongly correlated with each other, their alterations reflecting cell signaling and immunosuppression in cancer cells, and the occurrence of apoptotic events. Phosphatidylinositol alterations in tumors suggests the feasibility of cancer treatment approaches aiming at tuning the dysregulated PI3K/AKT pathway. Although MALDI MSI experiments have relatively lower lipid coverage when compared to LC-MS studies, their ability to generate highly specific lipid distribution maps within the tissues themselves provides invaluable biological information on OC progression mechanisms.

## Supporting Information

Mice tissue collection information, complete lipid feature list annotated with their putative IDs (Table S1), AUC for ROC curves, *p*-values for *t*-tests and log_2_FC of each group (tumor *vs*. control) (Table S2), mass spectrometry images of tissue sections not shown in Figure 7 (Figure S1) and mean mass spectra and PS/PA ratios measured at different laser powers (Figure S2).

## Notes

The authors declare no competing financial interest.

