## Supporting Information for "Ultrahigh Resolution Lipid Mass Spectrometry Imaging of High-Grade Serous Ovarian Cancer Mouse Models"

**Table of Contents**

Table S1. DKO, TKO and Control mice collection information	S1

Table S2. Complete lipid features annotated in the MSI experiments of each group	S2

Figure S1. Selected ion images and their mass errors of the remaining tissue sections as studied in the MSI experiments	S32

Figure S2. Investigations of the influence of laser powers on PS:PA peak areas	S36

**Table S1**. DKO, TKO and Control mice collection information

| **Mouse** | **Sample No.** | **Mouse ID** | **DOB** | **Collection Date** | **Age (Weeks)** | **Color** |
| --- | --- | --- | --- | --- | --- | --- |
| P2 | DKO-1 | 613 | 1/6/2020 | 8/4/2020 | 31 | Black |
| P6 | DKO-2 | 624 | 1/7/2020 | 8/4/2020 | 31 | White |
| P5 | DKO-3 | 623 | 1/7/2020 | 8/14/2020 | 32.5 | Black |
| P16 | DKO-4 | 657 | 1/6/2020 | 8/14/2020 | 32.5 | White |
| 6 | TKO-1 | 7529 | 4/16/2019 | 10/18/2019 | 27 | Black |
| 10 | TKO-2 | 8104 | 7/9/2019 | 1/17/2020 | 28 | Black |
| 1 | TKO-3 | 7320 | 3/24/2019 | 10/18/2019 | 30 | Black |
| 7 | TKO-4 | 8097 | 7/6/2019 | 12/20/2019 | 24 | White |
| 1 Ctrl | Control-1 | 7244 | 3/14/2019 | 12/20/2019 | 40 | Black |
| 4 Ctrl | Control-2 | 7531 | 4/12/2019 | 1/17/2020 | 40 | White |
| 5 Ctrl | Control-3 | 7532 | 4/12/2019 | 1/17/2020 | 40 | White |
| 7 Ctrl | Control-4 | 7781 | 5/17/2019 | 1/17/2020 | 35 | White |

**Table S2**. Lipid features annotated in the MSI experiments, their ROC curve AUC, *t*-test *p*-values, log_2_FC (Control/Tumor) / log_2_FC(Cyst/Tumor) of each group.

| Group | Putative Lipid ID | AUC | *p* Value | Log2FC |
| --- | --- | --- | --- | --- |
| DKO vs. Control | FA(22:4), Adrenic acid | 1 | 4.83E-16 | 1.7093 |
|  | PE(P-34:2), PE(O-34:3) | 1 | 2.29E-17 | -5.5049 |
|  | PE(P-34:0), PE(O-34:1), PC(P-31:0), PC(O-31:1) | 1 | 4.60E-18 | -3.9322 |
|  | PE(O-34:0), PC(O-31:0) | 1 | 2.29E-16 | -8.3672 |
|  | PE(P-36:2), PE(O-36:3), PC(P-33:2) | 1 | 8.13E-13 | -2.344 |
|  | PE(P-36:0), PE(O-36:1), PC(P-33:0), PC(O-33:1) | 1 | 1.84E-17 | -2.8731 |
|  | PE(34:0), PC(31:0) | 0.996753 | 3.70E-12 | -0.92901 |
|  | PE(36:2), PC(33:2) | 0.996753 | 1.89E-13 | -0.52299 |
|  | PA(39:4) | 0.993506 | 3.40E-15 | 1.7501 |
|  | PA(32:0) | 0.988636 | 9.89E-12 | -0.8747 |
|  | PA(O-34:1), PA(P-34:0) | 0.98539 | 2.03E-16 | -4.8485 |
|  | PA(O-36:3), PA(P-36:2) | 0.98539 | 3.62E-13 | -8.8829 |
|  | PE(40:4), PC(37:4) | 0.98539 | 1.03E-14 | 1.6138 |
|  | PA(40:5) | 0.982143 | 5.66E-16 | 1.3257 |
|  | PA(34:0) | 0.975649 | 7.55E-12 | -2.2783 |
|  | PE(32:0), PC(29:0) | 0.975649 | 2.89E-15 | -3.1891 |
|  | PI(36:3) | 0.969156 | 2.72E-13 | -1.4812 |
|  | PI(36:2) | 0.967532 | 1.29E-10 | -1.8802 |
|  | PE(P-38:5), PE(O-38:6) | 0.965909 | 2.18E-14 | -0.76943 |
|  | PG(34:0) | 0.964286 | 4.41E-14 | -0.78593 |
|  | PE(36:0), PC(33:0) | 0.961039 | 8.74E-10 | -1.1571 |
|  | FA(18:0), Stearic acid | 0.959416 | 1.15E-10 | 0.98209 |
|  | Cholesterol sulfate | 0.959416 | 2.75E-10 | 2.5626 |
|  | PI(O-38:5), PI(P-38:4) | 0.959416 | 8.92E-10 | -2.3193 |
|  | PI(38:4) | 0.959416 | 8.34E-10 | 0.78976 |
|  | PA(38:4) | 0.954545 | 4.21E-11 | 1.1466 |
|  | PE(38:4), PC(35:4) | 0.952922 | 1.58E-11 | 1.0687 |
|  | PE(P-38:6) | 0.951299 | 5.26E-12 | -1.4896 |
|  | PA(O-36:2), PA(P-36:1) | 0.948052 | 6.84E-09 | -3.9269 |
|  | PG(40:7) | 0.946429 | 1.46E-08 | -1.6281 |
|  | PA(O-38:6), PA(P-38:5) | 0.943182 | 1.66E-08 | -1.4761 |
|  | PE(P-36:1), PE(O-36:2), PC(P-33:1), PC(O-33:2) | 0.943182 | 3.80E-09 | -2.0975 |
|  | CPA(18:0) | 0.939935 | 3.19E-10 | 0.91566 |
|  | PA(33:0) | 0.939935 | 6.34E-10 | -2.461 |
|  | PE(39:4), PC(36:4) | 0.939935 | 7.36E-09 | 1.3619 |
|  | PA(32:1) | 0.935065 | 2.12E-09 | -2.0317 |
|  | PE(38:4(12OH)) | 0.935065 | 1.89E-07 | 2.123 |
|  | PA(O-36:4), PA(P-36:3) | 0.931818 | 1.35E-08 | -4.2971 |
|  | FA(22:5), DPA | 0.931818 | 7.06E-09 | 1.1962 |
|  | PE(P-16:0) | 0.928571 | 3.55E-10 | -0.72474 |
|  | SM(d35:1), PE-Cer(d38:1) | 0.928571 | 1.13E-06 | 1.2945 |
|  | PA(18:0), LPA(18:0) | 0.926948 | 7.21E-10 | 0.79416 |
|  | PA(34:2) | 0.926948 | 1.23E-08 | -0.46055 |
|  | PA(O-36:5), PA(P-36:4) | 0.923701 | 1.72E-07 | -0.98454 |
|  | PS(36:2) | 0.918831 | 1.09E-08 | -1.4784 |
|  | PIP(38:4) | 0.918831 | 1.59E-08 | 3.9529 |
|  | 1-Oleoylglycerophosphoinositol | 0.913961 | 3.44E-08 | -2.7805 |
|  | PA(37:4) | 0.913961 | 1.31E-07 | 1.0879 |
|  | PA(O-34:2), PA(P-34:1) | 0.912338 | 4.39E-08 | -4.2061 |
|  | PE(38:3), PC(35:3) | 0.907468 | 4.18E-10 | -0.59523 |
|  | PE(20:0), LPE(20:0), PC(17:0), PC(O-17:0) | 0.902597 | 3.25E-07 | 1.1583 |
|  | PE(34:2), PC(31:2) | 0.88474 | 5.27E-07 | -0.74558 |
|  | PE(P-36:4), PE(O-36:5) | 0.881494 | 1.25E-08 | -0.54532 |
|  | PE(P-38:1), PE(O-38:2), PC(P-35:1), PC(O-35:2) | 0.87987 | 3.65E-07 | -3.6924 |
|  | PE(P-42:6) | 0.87987 | 2.97E-07 | -1.6884 |
|  | PA(36:4) | 0.876623 | 4.77E-07 | 0.85808 |
|  | PS(44:8) | 0.875 | 1.49E-06 | 1.1958 |
|  | PA(36:3) | 0.87013 | 1.26E-06 | -0.3357 |
|  | PE(P-34:1), PE(O-34:2), PC(P-31:1) | 0.863636 | 6.11E-07 | -0.98933 |
|  | PS(44:3) | 0.863636 | 2.14E-05 | 3.2871 |
|  | PA(40:4) | 0.862013 | 9.21E-07 | 0.90513 |
|  | PE(P-40:7) | 0.862013 | 1.32E-07 | -1.8345 |
|  | PE(P-40:5), PE(O-40:6) | 0.858766 | 3.57E-06 | 0.58155 |
|  | PE(34:1), PC(31:1) | 0.855519 | 1.47E-06 | -0.74282 |
|  | PE-Cer(d36:1(2OH)) | 0.847403 | 3.88E-05 | 1.5343 |
|  | PE(P-38:0), PE(O-38:1), PC(P-35:0), PC(O-35:1) | 0.847403 | 5.90E-06 | -4.8231 |
|  | FA(20:4), Arachidonic acid | 0.845779 | 1.78E-06 | 0.78612 |
|  | PE(42:4), PC(39:4) | 0.845779 | 1.20E-06 | 1.0301 |
|  | PE(42:5), PC(39:5) | 0.844156 | 1.84E-06 | 0.76346 |
|  | PA(P-16:0) | 0.842532 | 0.000156 | -1.6359 |
|  | PA(36:2) | 0.840909 | 1.55E-06 | -0.36978 |
|  | PA(35:2) | 0.834416 | 1.79E-05 | -0.78697 |
|  | PG(36:0) | 0.832792 | 2.84E-05 | 0.55476 |
|  | FA(24:4) | 0.831169 | 8.88E-06 | 1.8985 |
|  | PI(39:4) | 0.829545 | 3.11E-05 | 0.98376 |
|  | PI(34:2) | 0.824675 | 2.75E-05 | -0.81995 |
|  | FA(18:2) | 0.821429 | 3.85E-05 | -0.36001 |
|  | PA(38:5) | 0.821429 | 3.29E-05 | 0.63166 |
|  | PI(40:7) | 0.816558 | 4.26E-05 | -4.1672 |
|  | PA(20:1) | 0.813312 | 5.67E-05 | 0.94417 |
|  | PE(38:1), PC(35:1) | 0.813312 | 5.09E-06 | 0.71483 |
|  | PS(34:1) | 0.811688 | 0.003199 | -1.917 |
|  | PG(44:8) | 0.808442 | 1.04E-07 | 0.42351 |
|  | PE(38:2), PC(35:2) | 0.805195 | 1.88E-05 | -0.47125 |
|  | SM(d41:1) | 0.805195 | 0.000382 | -0.92213 |
|  | PS(44:6) | 0.805195 | 0.000185 | -2.1359 |
|  | PA(P-38:6) | 0.803571 | 4.58E-05 | -3.2935 |
|  | PE(P-38:2) | 0.803571 | 1.42E-05 | -2.3983 |
|  | PGP(37:0) | 0.803571 | 1.08E-06 | 0.86882 |
|  | PE(36:4), PE(P-36:4), PC(33:4) | 0.800325 | 0.000165 | 0.74596 |
|  | PG(36:3) | 0.797078 | 3.15E-05 | -1.4702 |
|  | PS(42:4) | 0.795455 | 0.000385 | -1.6852 |
|  | PS(40:4) | 0.790584 | 0.000413 | 1.1983 |
|  | PI(40:6) | 0.788961 | 0.000252 | -0.41548 |
|  | PS(18:0) | 0.785714 | 0.000145 | 1.4577 |
|  | PA(38:3) | 0.784091 | 0.000665 | -0.21865 |
|  | PE(P-38:3), PE(O-38:4), PC(O-35:4) | 0.782468 | 4.18E-06 | 1.0614 |
|  | PE(39:7), PC(36:7) | 0.780844 | 0.000779 | -4.4253 |
|  | PE(42:6), PC(39:6) | 0.780844 | 0.000124 | 0.63271 |
|  | PI(34:0) | 0.772727 | 0.00021 | 1.6052 |
|  | PA(20:4) | 0.771104 | 0.000944 | 0.80886 |
|  | PE(O-18:0), PC(O-15:0) | 0.771104 | 2.10E-05 | 1.2433 |
|  | PE(38:5), PC(35:5) | 0.767857 | 0.000206 | 0.65542 |
|  | FA(16:0), Palmitic acid | 0.766234 | 0.000595 | -0.22304 |
|  | PG(36:4) | 0.761364 | 0.001474 | -1.5681 |
|  | PI(38:3) | 0.761364 | 0.000341 | -0.56662 |
|  | PE(36:3), PC(33:3) | 0.75974 | 0.004386 | -0.30678 |
|  | CerP(d44:2) | 0.758117 | 0.001347 | -1.5769 |
|  | PE(40:5), PC(37:5) | 0.75 | 0.000584 | 0.50064 |
|  | CerP(d36:1) | 0.748377 | 0.009613 | 0.62626 |
|  | PS(40:5) | 0.748377 | 0.003556 | 1.0311 |
|  | PE(39:5), PC(36:5) | 0.741883 | 0.002091 | 0.74391 |
|  | PA(38:6) | 0.741883 | 0.00109 | -0.18919 |
|  | CerP(d42:1) | 0.74026 | 0.003082 | -1.7048 |
|  | PE(P-36:3), PE(O-36:4) | 0.738636 | 0.027522 | 0.48214 |
|  | PE(P-40:6) | 0.738636 | 0.00392 | -0.296 |
|  | FA(20:2) | 0.73539 | 0.031773 | -0.11646 |
|  | PE(44:10) | 0.728896 | 0.000112 | 1.3276 |
|  | PI(O-36:4), PI(P-36:3) | 0.728896 | 0.000352 | -3.6849 |
|  | SM(d33:2), PE-Cer(d36:2) | 0.727273 | 0.019236 | 1.4368 |
|  | PE(22:4), LPE(22:4) | 0.719156 | 0.002205 | 0.87768 |
|  | PS(38:3) | 0.717532 | 0.025022 | -0.70447 |
|  | PG(36:1) | 0.714286 | 0.029857 | -0.28131 |
|  | PA(34:1) | 0.709416 | 0.026991 | -0.29603 |
|  | PS(38:4) | 0.709416 | 0.010883 | 0.83184 |
|  | PI(38:5) | 0.709416 | 9.35E-05 | -0.90046 |
|  | CPA(18:2) | 0.707792 | 0.050433 | -0.00572 |
|  | PE(20:2), LPE(20:2), PC(17:2) | 0.706169 | 0.03858 | -0.9014 |
|  | PE(18:0), LPE(18:0), PC(15:0) | 0.704545 | 0.023386 | 0.45419 |
|  | PG(36:2) | 0.704545 | 0.016128 | -2.7614 |
|  | PI(20:4) | 0.694805 | 0.005429 | 0.76804 |
|  | PS(44:5) | 0.694805 | 0.054655 | -0.54935 |
|  | Cer(d42:2) | 0.689935 | 0.040027 | -2.2504 |
|  | DHAP(18:0) | 0.688312 | 0.052468 | -0.48021 |
|  | PG(44:10) | 0.681818 | 0.010314 | 0.34533 |
|  | PA(O-40:6), PA(P-40:5) | 0.678571 | 0.009868 | -0.84957 |
|  | CerP(d34:0) | 0.676948 | 0.016422 | -1.8744 |
|  | PI(40:3) | 0.676948 | 0.188413 | -2.7229 |
|  | PA(18:1) | 0.675325 | 0.023023 | -0.25296 |
|  | 1-hexadecanyl-2-(8-[3]-ladderane-octanyl)-sn-glycerophosphocholine | 0.675325 | 0.007599 | -0.57106 |
|  | PE(42:8), PC(39:8) | 0.670455 | 0.005295 | 0.71429 |
|  | PI(40:5) | 0.668831 | 0.059188 | -0.05176 |
|  | PA(36:1) | 0.667208 | 0.016918 | 0.40344 |
|  | LPE(22:6) | 0.665584 | 0.376738 | -0.96803 |
|  | PG(38:5) | 0.663961 | 0.003469 | -1.747 |
|  | PE(P-40:3), PE(O-40:4), PC(O-37:4) | 0.662338 | 0.001989 | 0.56712 |
|  | PI(36:1) | 0.660714 | 0.037527 | -0.63028 |
|  | PS(42:1) | 0.655844 | 0.12756 | -1.9712 |
|  | PE(P-20:0), PC(P-17:0) | 0.652597 | 0.101473 | -0.66257 |
|  | PE(37:5), PC(34:5) | 0.652597 | 0.238747 | -0.40424 |
|  | PE(40:8) | 0.652597 | 0.004729 | 1.4752 |
|  | PG(42:8) | 0.652597 | 0.014917 | -0.70406 |
|  | FA(20:3) | 0.650974 | 0.210988 | 0.015006 |
|  | PE(P-40:4), PE(O-40:5) | 0.646104 | 0.067864 | 0.40664 |
|  | Cer(d34:1) | 0.642857 | 0.07562 | -2.2293 |
|  | PA(37:2) | 0.642857 | 0.503463 | -0.43385 |
|  | PA(40:7) | 0.642857 | 0.19045 | -0.46654 |
|  | PE(36:1), PC(33:1) | 0.63961 | 0.196934 | -0.22768 |
|  | PE(38:6), PC(35:6) | 0.637987 | 0.646 | -0.06238 |
|  | PG(44:11) | 0.636364 | 0.021566 | 0.65403 |
|  | PA(O-38:5), PA(P-38:4) | 0.63474 | 0.004255 | -0.93301 |
|  | PE(40:6), PC(37:6) | 0.63474 | 0.914704 | 0.17133 |
|  | FA(18:1), Oleic acid | 0.633117 | 0.115773 | -0.37885 |
|  | PE(37:2), PC(34:2) | 0.631494 | 0.155406 | -0.45882 |
|  | PI(34:1) | 0.631494 | 0.156878 | -0.30061 |
|  | PE(39:6), PC(36:6) | 0.628247 | 0.204443 | -0.67701 |
|  | PS(44:4) | 0.626623 | 0.380256 | 0.68773 |
|  | PE(20:1), LPE(20:1), PC(17:1) | 0.625 | 0.631986 | -0.74135 |
|  | PI(O-40:4), PI(P-40:3) | 0.625 | 0.19554 | 0.25559 |
|  | PI(18:0) | 0.623377 | 0.17534 | 0.29028 |
|  | CPA(18:1) | 0.621753 | 0.129902 | -0.41565 |
|  | PS(40:1) | 0.621753 | 0.136754 | 0.33106 |
|  | SM(d41:2) | 0.62013 | 0.024905 | -0.43837 |
|  | PI(40:4) | 0.618506 | 0.142493 | 0.46683 |
|  | PG(42:10) | 0.616883 | 0.036359 | -1.6773 |
|  | PG(40:5) | 0.61526 | 0.698195 | -1.2142 |
|  | PA(O-38:4), PA(P-38:3) | 0.613636 | 0.064775 | -0.197 |
|  | PE(42:7), PC(39:7) | 0.612013 | 0.467635 | -0.87565 |
|  | SM(d33:0) | 0.608766 | 0.045307 | -1.0665 |
|  | PG(38:6) | 0.608766 | 0.02746 | -2.3549 |
|  | PE(O-42:6) | 0.608766 | 0.081673 | -0.9805 |
|  | PG(40:8) | 0.608766 | 0.041491 | -1.8302 |
|  | CerP(d42:2) | 0.607143 | 0.026453 | -0.38119 |
|  | SM(d37:1), PE-Cer(d40:1) | 0.607143 | 0.05071 | -1.1406 |
|  | PA(42:7) | 0.607143 | 0.006399 | 0.38341 |
|  | PG(44:12) | 0.605519 | 0.279508 | 0.021607 |
|  | PA(37:3) | 0.603896 | 0.528426 | -1.5592 |
|  | PA(38:2) | 0.602273 | 0.006985 | 0.36955 |
|  | PG(42:9) | 0.600649 | 0.173551 | -0.65505 |
|  | PE(40:3), PC(37:3) | 0.599026 | 0.031305 | 0.31068 |
|  | PE(40:7), PC(37:7) | 0.595779 | 0.867002 | -0.3144 |
|  | PI(37:4) | 0.592532 | 0.078068 | 0.30943 |
|  | FA(20:1) | 0.592532 | 0.630354 | -1.2153 |
|  | SM(d39:1) | 0.582792 | 0.593632 | -0.11492 |
|  | FA(22:6), DHA | 0.581169 | 0.297038 | -0.32013 |
|  | CerP(d34:1) | 0.581169 | 0.218767 | 0.36102 |
|  | SM(d39:2) | 0.579545 | 0.781153 | -0.76814 |
|  | PI(38:6) | 0.579545 | 0.852031 | -0.004 |
|  | PE(44:8) | 0.577922 | 0.126564 | 0.44523 |
|  | PA(35:1) | 0.571429 | 0.374064 | -0.20839 |
|  | PA(O-40:5), PA(P-40:4) | 0.568182 | 0.532843 | -0.54754 |
|  | PI(O-38:4), PI(P-38:3) | 0.568182 | 0.565261 | -0.63378 |
|  | PE(P-38:4), PE(O-38:5) | 0.566558 | 0.166313 | 0.4024 |
|  | PE(40:2), PC(37:2) | 0.566558 | 0.74482 | -1.017 |
|  | 22:0-Glc-Sitosterol | 0.560065 | 0.561254 | 0.74031 |
|  | PS(P-38:4), PS(O-38:5) | 0.556818 | 0.992949 | -0.27454 |
|  | PA(40:6) | 0.555195 | 0.813797 | 0.15313 |
|  | PI(36:4) | 0.551948 | 0.008031 | 0.41712 |
|  | PC(20:4) | 0.547078 | 0.11843 | -0.35676 |
|  | PG(38:4) | 0.540584 | 0.340024 | -1.0996 |
|  | CPA(16:0) | 0.538961 | 0.368955 | 0.24687 |
|  | SM(d33:1), PE-Cer(d36:1) | 0.538961 | 0.824384 | 0.16581 |
|  | PE(20:4), LPE(20:4) | 0.535714 | 0.734022 | 0.33219 |
|  | PG(40:6) | 0.534091 | 0.136686 | -1.2187 |
|  | PE(18:1), LPE(18:1), PC(15:1) | 0.532468 | 0.178961 | -0.1117 |
|  | PC(dO-34:4) | 0.530844 | 0.215174 | -1.0221 |
|  | PG(34:1) | 0.529221 | 0.970148 | 0.17622 |
|  | PC(36:2) | 0.529221 | 0.604716 | -1.5058 |
|  | PS(40:6) | 0.525974 | 0.732488 | 0.10293 |
|  | PE(P-18:0), PE(O-18:1), PC(P-15:0) | 0.524351 | 0.699744 | 0.40657 |
|  | PE(16:0), LPE(16:0), PC(13:0) | 0.522727 | 0.155765 | 0.22735 |
|  | PI(16:0) | 0.522727 | 0.025761 | 0.10774 |
|  | PI(38:2) | 0.516234 | 0.857171 | 0.084205 |
|  | PS(36:1) | 0.512987 | 0.427946 | 0.25032 |
|  | PE(P-42:4) | 0.508117 | 0.753655 | -1.2813 |
|  | PA(16:0), LPA(16:0) | 0.50487 | 0.757002 | 0.19883 |
|  | PS(38:1) | 0.50487 | 0.936664 | 0.4237 |
|  | CerP(d40:1) | 0.501623 | 0.084388 | -0.21685 |
| TKO vs. Control | FA(20:1) | 1 | 5.89E-13 | -1.5602 |
|  | CPA(18:1) | 1 | 4.12E-18 | -0.70539 |
|  | Cholesterol sulfate | 1 | 4.17E-14 | 4.6785 |
|  | PE(O-18:0), PC(O-15:0) | 1 | 6.18E-18 | 3.9563 |
|  | PE(20:0), LPE(20:0), PC(17:0), PC(O-17:0) | 1 | 1.82E-16 | 1.9361 |
|  | 1-Oleoylglycerophosphoinositol | 1 | 3.78E-16 | -3.0022 |
|  | CerP(d36:1) | 1 | 9.10E-13 | 2.0604 |
|  | PA(32:1) | 1 | 3.27E-15 | -2.7244 |
|  | PA(33:0) | 1 | 2.15E-15 | -3.3601 |
|  | PE(32:0), PC(29:0) | 1 | 1.51E-20 | -3.8201 |
|  | CerP(d40:1) | 1 | 1.86E-15 | 3.425 |
|  | PE(P-34:0), PE(O-34:1), PC(P-31:0), PC(O-31:1) | 1 | 2.96E-11 | -2.6046 |
|  | PA(O-38:4), PA(P-38:3) | 1 | 1.85E-14 | 3.0878 |
|  | SM(d35:1), PE-Cer(d38:1) | 1 | 3.06E-16 | 3.9546 |
|  | PE(P-36:4), PE(O-36:5) | 1 | 1.35E-14 | -0.76387 |
|  | PA(39:4) | 1 | 4.42E-14 | 1.3116 |
|  | PE(P-38:5), PE(O-38:6) | 1 | 1.33E-17 | -0.73387 |
|  | PG(34:0) | 1 | 2.21E-16 | -0.749 |
|  | PE(P-38:3), PE(O-38:4), PC(O-35:4) | 1 | 4.75E-22 | 2.4395 |
|  | PE(38:4), PC(35:4) | 1 | 3.06E-12 | 0.85093 |
|  | PE(P-40:3), PE(O-40:4), PC(O-37:4) | 1 | 6.09E-15 | 1.738 |
|  | PE(40:4), PC(37:4) | 1 | 7.61E-19 | 1.6506 |
|  | PI(38:5) | 1 | 4.92E-19 | -1.1952 |
|  | PI(40:6) | 1 | 4.62E-11 | -1.3373 |
|  | PE(P-38:6) | 0.997768 | 4.34E-15 | -1.061 |
|  | PE(P-40:7) | 0.997768 | 8.78E-18 | -1.1096 |
|  | PG(36:2) | 0.997768 | 4.94E-13 | -1.7308 |
|  | FA(20:2) | 0.995536 | 2.96E-13 | -0.69556 |
|  | PA(34:0) | 0.995536 | 5.15E-13 | -2.9416 |
|  | PA(38:4) | 0.995536 | 4.75E-12 | 1.0379 |
|  | PI(40:5) | 0.995536 | 8.28E-12 | -0.9108 |
|  | PE(36:0), PC(33:0) | 0.993304 | 2.60E-11 | -1.3872 |
|  | PE(38:1), PC(35:1) | 0.993304 | 1.66E-14 | 1.398 |
|  | PE(34:1), PC(31:1) | 0.991071 | 1.20E-13 | -0.86898 |
|  | PE(40:7), PC(37:7) | 0.991071 | 5.90E-15 | -0.56272 |
|  | PG(40:7) | 0.991071 | 3.07E-12 | -1.3683 |
|  | PI(40:3) | 0.991071 | 2.36E-11 | -4.7706 |
|  | FA(18:0), Stearic acid | 0.988839 | 7.47E-13 | 0.76319 |
|  | PE(P-36:3), PE(O-36:4) | 0.988839 | 5.55E-13 | 1.3688 |
|  | PA(40:5) | 0.988839 | 8.81E-12 | 1.0166 |
|  | PE(42:7), PC(39:7) | 0.986607 | 2.42E-09 | -0.57845 |
|  | PG(40:6) | 0.986607 | 2.89E-12 | -1.1165 |
|  | PI(36:3) | 0.986607 | 2.81E-13 | -1.4468 |
|  | PE(O-34:0), PC(O-31:0) | 0.984375 | 3.49E-10 | -5.0458 |
|  | PI(40:7) | 0.984375 | 3.26E-13 | -5.2654 |
|  | PA(40:7) | 0.982143 | 1.46E-07 | -0.51268 |
|  | PE(39:4), PC(36:4) | 0.979911 | 1.00E-09 | 1.3604 |
|  | SM(d39:1) | 0.977679 | 3.53E-10 | 1.5546 |
|  | PE(34:2), PC(31:2) | 0.975446 | 2.70E-09 | -1.0412 |
|  | PE(42:4), PC(39:4) | 0.975446 | 1.27E-10 | 0.92787 |
|  | PS(34:1) | 0.973214 | 4.74E-09 | -2.5696 |
|  | CerP(d34:1) | 0.970982 | 3.50E-11 | 0.98453 |
|  | PE(34:0), PC(31:0) | 0.970982 | 1.15E-12 | -1.0388 |
|  | PI(38:3) | 0.970982 | 1.12E-09 | -0.99237 |
|  | PA(18:1) | 0.96875 | 2.73E-11 | -0.41192 |
|  | PA(35:2) | 0.96875 | 6.68E-08 | -1.3248 |
|  | PI(36:2) | 0.96875 | 1.41E-09 | -0.94328 |
|  | PS(42:1) | 0.96875 | 8.36E-10 | -6.0292 |
|  | PA(20:1) | 0.966518 | 9.33E-08 | 1.3417 |
|  | PA(O-38:6), PA(P-38:5) | 0.966518 | 1.24E-06 | -1.1314 |
|  | Cer(d42:2) | 0.964286 | 5.24E-09 | -4.3199 |
|  | PA(32:0) | 0.964286 | 4.60E-12 | -1.0636 |
|  | PA(36:3) | 0.962054 | 5.24E-09 | -0.62589 |
|  | PA(36:2) | 0.962054 | 8.57E-09 | -0.24032 |
|  | PE-Cer(d36:1(2OH)) | 0.962054 | 5.08E-10 | 2.7357 |
|  | PG(38:5) | 0.962054 | 2.05E-10 | -0.93619 |
|  | Cer(d34:1) | 0.959821 | 9.93E-09 | -4.3193 |
|  | PE(38:3), PC(35:3) | 0.959821 | 1.42E-09 | -0.82186 |
|  | PE(39:7), PC(36:7) | 0.959821 | 6.05E-09 | -4.4739 |
|  | PA(34:2) | 0.957589 | 3.26E-09 | -0.53355 |
|  | PA(O-36:5), PA(P-36:4) | 0.955357 | 7.18E-07 | -0.94584 |
|  | PA(35:1) | 0.955357 | 4.75E-10 | -0.46835 |
|  | SM(d33:1), PE-Cer(d36:1) | 0.950893 | 1.36E-09 | 1.0094 |
|  | PE(20:1), LPE(20:1), PC(17:1) | 0.948661 | 5.25E-06 | -0.53845 |
|  | PE(P-34:2), PE(O-34:3) | 0.948661 | 4.15E-08 | -3.1289 |
|  | PA(37:2) | 0.946429 | 4.08E-07 | -0.56894 |
|  | PS(36:2) | 0.946429 | 1.20E-08 | -1.3202 |
|  | CPA(18:2) | 0.944196 | 4.01E-08 | -0.48316 |
|  | PE(P-16:0) | 0.944196 | 3.39E-09 | -0.82344 |
|  | PA(18:0), LPA(18:0) | 0.933036 | 2.18E-08 | 0.68716 |
|  | PE(36:3), PC(33:3) | 0.933036 | 6.35E-07 | -1.0339 |
|  | FA(16:0), Palmitic acid | 0.928571 | 1.84E-08 | -0.41022 |
|  | PA(37:3) | 0.928571 | 5.92E-07 | -2.3178 |
|  | LPE(22:6) | 0.926339 | 0.000114 | -1.1057 |
|  | PE(36:2), PC(33:2) | 0.926339 | 9.78E-08 | -0.51312 |
|  | PG(38:6) | 0.912946 | 3.75E-07 | -0.74784 |
|  | PA(O-34:1), PA(P-34:0) | 0.908482 | 1.41E-06 | -2.1999 |
|  | PI(40:4) | 0.908482 | 2.25E-06 | -0.35373 |
|  | PA(O-40:5), PA(P-40:4) | 0.90625 | 7.58E-07 | 1.5894 |
|  | CPA(18:0) | 0.904018 | 6.33E-07 | 0.612 |
|  | PG(36:3) | 0.904018 | 1.80E-05 | -1.0146 |
|  | PG(42:9) | 0.904018 | 6.40E-07 | -0.83862 |
|  | PE(P-34:1), PE(O-34:2), PC(P-31:1) | 0.901786 | 6.82E-06 | -0.45604 |
|  | PG(40:5) | 0.901786 | 8.06E-08 | -1.2215 |
|  | PE(P-36:0), PE(O-36:1), PC(P-33:0), PC(O-33:1) | 0.897321 | 1.71E-06 | -0.97578 |
|  | PS(44:3) | 0.897321 | 4.87E-07 | 4.2643 |
|  | PI(O-38:5), PI(P-38:4) | 0.895089 | 5.63E-06 | -0.93863 |
|  | PC(dO-34:4) | 0.888393 | 2.96E-05 | 1.8347 |
|  | PE(P-40:6) | 0.888393 | 2.64E-06 | -0.13499 |
|  | PG(36:1) | 0.881696 | 4.57E-06 | -0.1476 |
|  | PE(37:5), PC(34:5) | 0.879464 | 3.19E-06 | -1.228 |
|  | PA(34:1) | 0.877232 | 1.68E-07 | -0.29261 |
|  | PE(18:1), LPE(18:1), PC(15:1) | 0.875 | 5.11E-05 | -0.31286 |
|  | PG(42:10) | 0.872768 | 8.30E-05 | -0.63821 |
|  | PG(40:8) | 0.870536 | 9.11E-06 | -0.57972 |
|  | PA(P-38:6) | 0.868304 | 2.93E-05 | -2.2913 |
|  | PE(P-40:5), PE(O-40:6) | 0.863839 | 5.58E-06 | 0.69432 |
|  | PE(P-40:4), PE(O-40:5) | 0.863839 | 3.89E-05 | 0.76451 |
|  | SM(d37:1), PE-Cer(d40:1) | 0.861607 | 7.92E-08 | 1.785 |
|  | PE(42:6), PC(39:6) | 0.861607 | 9.18E-06 | 0.84117 |
|  | PE(20:2), LPE(20:2), PC(17:2) | 0.861607 | 0.000271 | -1.1553 |
|  | PG(36:4) | 0.861607 | 0.000193 | -0.7622 |
|  | PI(38:6) | 0.859375 | 0.000239 | -0.643 |
|  | PS(38:3) | 0.857143 | 0.0001 | -0.98485 |
|  | FA(20:4), Arachidonic acid | 0.857143 | 0.000104 | 0.64391 |
|  | PA(38:3) | 0.852679 | 1.23E-05 | -0.22053 |
|  | SM(d39:2) | 0.852679 | 6.15E-05 | 1.8756 |
|  | PE(39:6), PC(36:6) | 0.852679 | 3.68E-05 | -1.2949 |
|  | PE(P-42:4) | 0.852679 | 1.04E-05 | 1.4485 |
|  | PA(40:6) | 0.845982 | 2.38E-05 | -0.12107 |
|  | PE(44:10) | 0.84375 | 0.000373 | 2.77 |
|  | PS(18:0) | 0.839286 | 6.51E-05 | 1.8206 |
|  | PA(P-16:0) | 0.834821 | 0.000891 | -1.5944 |
|  | PA(O-38:5), PA(P-38:4) | 0.832589 | 0.00034 | -0.19484 |
|  | PG(36:0) | 0.828125 | 6.14E-05 | 0.65357 |
|  | PI(O-38:4), PI(P-38:3) | 0.828125 | 0.000262 | 1.0749 |
|  | FA(22:4), Adrenic acid | 0.825893 | 0.000118 | 0.67852 |
|  | PS(P-38:4), PS(O-38:5) | 0.821429 | 0.00141 | -1.2761 |
|  | CerP(d42:2) | 0.819196 | 0.000135 | 0.7664 |
|  | PE(20:4), LPE(20:4) | 0.816964 | 0.000571 | -0.3773 |
|  | PS(44:8) | 0.814732 | 9.52E-05 | 0.84625 |
|  | PA(O-36:3), PA(P-36:2) | 0.8125 | 6.53E-05 | -4.5919 |
|  | PA(36:4) | 0.8125 | 0.000237 | 0.66045 |
|  | PI(38:4) | 0.8125 | 0.000435 | 0.50637 |
|  | PA(38:6) | 0.810268 | 0.000137 | -0.10906 |
|  | FA(20:3) | 0.803571 | 8.61E-05 | -0.32614 |
|  | PI(39:4) | 0.803571 | 0.00076 | 0.81394 |
|  | PE(40:2), PC(37:2) | 0.794643 | 0.000504 | -0.49201 |
|  | PG(44:8) | 0.792411 | 0.000744 | 1.08 |
|  | PG(44:12) | 0.790179 | 0.000313 | 0.83422 |
|  | SM(d41:2) | 0.787946 | 0.000134 | 0.82273 |
|  | PE(O-42:6) | 0.787946 | 0.000322 | 1.1463 |
|  | PE(44:8) | 0.779018 | 0.001751 | -0.63714 |
|  | FA(18:1), Oleic acid | 0.772321 | 0.000645 | -0.24274 |
|  | PS(40:6) | 0.765625 | 0.002987 | -0.34519 |
|  | PS(44:4) | 0.765625 | 0.074652 | 1.0341 |
|  | PA(40:4) | 0.756696 | 0.008511 | 0.48434 |
|  | PS(38:1) | 0.754464 | 0.003804 | -1.1928 |
|  | PI(34:2) | 0.75 | 0.012208 | -0.4448 |
|  | PE(40:6), PC(37:6) | 0.741071 | 0.004857 | 0.08974 |
|  | PS(40:1) | 0.738839 | 0.003828 | -1.6099 |
|  | FA(24:4) | 0.734375 | 0.011114 | -0.96461 |
|  | PE(36:1), PC(33:1) | 0.732143 | 0.001051 | 0.006568 |
|  | CPA(16:0) | 0.729911 | 0.006788 | 0.042761 |
|  | PA(36:1) | 0.723214 | 0.019893 | 0.47842 |
|  | PS(38:4) | 0.723214 | 0.118995 | 0.58957 |
|  | PS(44:6) | 0.723214 | 0.072038 | -1.0182 |
|  | PE(P-20:0), PC(P-17:0) | 0.71875 | 0.029509 | 2.9969 |
|  | PG(44:10) | 0.707589 | 0.010406 | -0.24983 |
|  | PG(34:1) | 0.705357 | 0.074437 | -0.05166 |
|  | PE(38:2), PC(35:2) | 0.698661 | 0.031746 | -0.0346 |
|  | PE(16:0), LPE(16:0), PC(13:0) | 0.696429 | 0.023616 | -0.05465 |
|  | PA(38:2) | 0.696429 | 0.032389 | 0.48106 |
|  | PG(38:4) | 0.691964 | 0.099819 | -0.3433 |
|  | PE(42:5), PC(39:5) | 0.691964 | 0.019181 | 0.52359 |
|  | PE(P-36:2), PE(O-36:3), PC(P-33:2) | 0.6875 | 0.044079 | -0.54702 |
|  | PS(40:5) | 0.685268 | 0.05506 | -0.18492 |
|  | PI(16:0) | 0.683036 | 0.04621 | -0.39656 |
|  | PE(18:0), LPE(18:0), PC(15:0) | 0.680804 | 0.028669 | 0.49519 |
|  | PE(38:6), PC(35:6) | 0.676339 | 0.027001 | 0.030208 |
|  | PA(16:0), LPA(16:0) | 0.671875 | 0.099894 | 0.10761 |
|  | PA(O-36:4), PA(P-36:3) | 0.671875 | 0.052761 | -0.61634 |
|  | PI(37:4) | 0.669643 | 0.806477 | -0.01939 |
|  | PG(42:8) | 0.662946 | 0.047927 | -0.17527 |
|  | PI(18:0) | 0.658482 | 0.121888 | 0.019356 |
|  | SM(d33:2), PE-Cer(d36:2) | 0.65625 | 0.058619 | 0.87659 |
|  | PE(40:5), PC(37:5) | 0.65625 | 0.024146 | 0.083013 |
|  | PA(O-34:2), PA(P-34:1) | 0.651786 | 0.038662 | -0.11896 |
|  | 1-hexadecanyl-2-(8-[3]-ladderane-octanyl)-sn-glycerophosphocholine | 0.649554 | 0.038732 | 0.58824 |
|  | PC(36:2) | 0.649554 | 0.110212 | -0.08989 |
|  | PGP(37:0) | 0.647321 | 0.120166 | -0.42849 |
|  | FA(18:2) | 0.645089 | 0.122294 | -0.26767 |
|  | FA(22:5), DPA | 0.640625 | 0.057645 | 0.54956 |
|  | PI(O-36:4), PI(P-36:3) | 0.640625 | 0.182639 | -0.60195 |
|  | PA(O-36:2), PA(P-36:1) | 0.638393 | 0.149244 | -0.67879 |
|  | PA(20:4) | 0.638393 | 0.073393 | 0.47694 |
|  | PS(42:4) | 0.636161 | 0.205723 | -0.63291 |
|  | PE(P-38:2) | 0.633929 | 0.027822 | 0.12117 |
|  | PI(38:2) | 0.631696 | 0.051873 | -1.2209 |
|  | PIP(38:4) | 0.629464 | 0.165989 | 1.0692 |
|  | PE(22:4), LPE(22:4) | 0.620536 | 0.181937 | 0.010396 |
|  | SM(d33:0) | 0.618304 | 0.254447 | 0.56403 |
|  | PI(34:1) | 0.618304 | 0.071971 | 0.036698 |
|  | PA(O-40:6), PA(P-40:5) | 0.613839 | 0.301599 | -0.02454 |
|  | PA(42:7) | 0.604911 | 0.528718 | 0.41798 |
|  | PG(44:11) | 0.602679 | 0.269251 | 0.038021 |
|  | PS(44:5) | 0.600446 | 0.884768 | -0.13562 |
|  | SM(d41:1) | 0.598214 | 0.184706 | 0.2779 |
|  | PI(34:0) | 0.598214 | 0.198524 | -0.20379 |
|  | PC(20:4) | 0.595982 | 0.660898 | -0.59892 |
|  | PE(38:5), PC(35:5) | 0.595982 | 0.500536 | 0.12266 |
|  | DHAP(18:0) | 0.591518 | 0.549306 | 0.11059 |
|  | PI(O-40:4), PI(P-40:3) | 0.587054 | 0.445076 | 0.72435 |
|  | 22:0-Glc-Sitosterol | 0.580357 | 0.37236 | -0.65146 |
|  | PE(37:2), PC(34:2) | 0.578125 | 0.180882 | 0.016885 |
|  | FA(22:6), DHA | 0.571429 | 0.594048 | 0.12605 |
|  | PE(36:4), PE(P-36:4), PC(33:4) | 0.571429 | 0.31757 | 0.31852 |
|  | PE(38:4(12OH)) | 0.569196 | 0.297936 | 0.18556 |
|  | PE(P-42:6) | 0.569196 | 0.409389 | -0.05514 |
|  | PI(20:4) | 0.566964 | 0.226517 | 0.64889 |
|  | PS(40:4) | 0.566964 | 0.788927 | 0.28249 |
|  | PE(P-38:1), PE(O-38:2), PC(P-35:1), PC(O-35:2) | 0.564732 | 0.206511 | -0.44475 |
|  | PE(40:3), PC(37:3) | 0.560268 | 0.276493 | 0.28695 |
|  | PA(38:5) | 0.555804 | 0.693215 | 0.21672 |
|  | PE(P-18:0), PE(O-18:1), PC(P-15:0) | 0.553571 | 0.920948 | 0.36586 |
|  | PE(P-38:0), PE(O-38:1), PC(P-35:0), PC(O-35:1) | 0.551339 | 0.587505 | 0.59099 |
|  | CerP(d42:1) | 0.549107 | 0.361195 | 0.28298 |
|  | PE(P-36:1), PE(O-36:2), PC(P-33:1), PC(O-33:2) | 0.546875 | 0.418947 | 0.16814 |
|  | PS(36:1) | 0.544643 | 0.608013 | 0.17848 |
|  | PI(36:1) | 0.535714 | 0.928134 | 0.73125 |
|  | CerP(d34:0) | 0.529018 | 0.569255 | 0.77186 |
|  | PE(39:5), PC(36:5) | 0.529018 | 0.724794 | -0.07908 |
|  | PE(40:8) | 0.522321 | 0.872759 | 0.11336 |
|  | PI(36:4) | 0.522321 | 0.896195 | 0.18707 |
|  | PE(P-38:4), PE(O-38:5) | 0.517857 | 0.678435 | 0.19333 |
|  | PE(42:8), PC(39:8) | 0.508929 | 0.541112 | 0.42632 |
|  | CerP(d44:2) | 0.506696 | 0.928934 | -0.19224 |
|  | PA(37:4) | 0.504464 | 0.862729 | 0.25968 |
|  | CPA(16:0) | 1 | 5.91E-13 | -6.4319 |
|  | PE(16:0), LPE(16:0), PC(13:0) | 1 | 1.13E-10 | -4.4766 |
|  | PE(P-18:0), PE(O-18:1), PC(P-15:0) | 1 | 3.98E-09 | 0.66634 |
|  | CerP(d34:0) | 1 | 1.80E-05 | 2.3628 |
|  | PA(32:1) | 1 | 3.03E-15 | -7.9335 |
|  | PA(O-34:1), PA(P-34:0) | 1 | 3.36E-07 | 1.8563 |
|  | PA(O-36:2), PA(P-36:1) | 1 | 1.24E-06 | 2.0254 |
|  | SM(d33:0) | 1 | 2.55E-06 | 2.0589 |
|  | PE(P-34:0), PE(O-34:1), PC(P-31:0), PC(O-31:1) | 1 | 1.82E-08 | 1.1311 |
|  | PE(O-34:0), PC(O-31:0) | 1 | 1.14E-06 | 2.4498 |
|  | PA(37:4) | 1 | 2.69E-13 | -5.3775 |
|  | PA(38:6) | 1 | 2.59E-14 | -3.9602 |
|  | PA(38:5) | 1 | 1.87E-12 | -2.44 |
|  | CerP(d42:2) | 1 | 2.04E-08 | 1.4674 |
|  | CerP(d42:1) | 1 | 6.42E-07 | 2.2107 |
|  | PE(P-36:0), PE(O-36:1), PC(P-33:0), PC(O-33:1) | 1 | 6.46E-10 | 1.6251 |
|  | PE(36:4), PE(P-36:4), PC(33:4) | 1 | 1.74E-10 | -2.0751 |
|  | PA(40:7) | 1 | 1.72E-14 | -6.1883 |
|  | PA(40:6) | 1 | 1.45E-11 | -2.7903 |
|  | PA(40:5) | 1 | 5.08E-11 | -2.1255 |
|  | PE(P-38:1), PE(O-38:2), PC(P-35:1), PC(O-35:2) | 1 | 1.05E-06 | 2.2636 |
|  | PE(38:5), PC(35:5) | 1 | 6.76E-13 | -2.8946 |
|  | PA(42:7) | 1 | 9.56E-12 | -8.2659 |
|  | PE(40:6), PC(37:6) | 1 | 2.13E-14 | -2.6641 |
|  | PE(40:5), PC(37:5) | 1 | 2.78E-16 | -2.4119 |
|  | PE(40:3), PC(37:3) | 1 | 6.85E-13 | -3.1477 |
|  | PE(42:7), PC(39:7) | 1 | 1.29E-12 | -7.8373 |
|  | PE(42:6), PC(39:6) | 1 | 3.03E-15 | -3.6888 |
|  | PE(42:5), PC(39:5) | 1 | 4.09E-16 | -5.1175 |
|  | PA(O-36:3), PA(P-36:2) | 0.996212 | 1.31E-05 | 2.4165 |
|  | SM(d33:1), PE-Cer(d36:1) | 0.996212 | 2.86E-08 | 1.4466 |
|  | PA(39:4) | 0.996212 | 5.51E-12 | -3.9289 |
|  | PE(P-38:2) | 0.996212 | 3.05E-07 | 2.1881 |
|  | FA(22:6), DHA | 0.992424 | 3.32E-07 | -2.8756 |
|  | PA(O-34:2), PA(P-34:1) | 0.992424 | 5.62E-06 | 2.3176 |
|  | PE(P-36:1), PE(O-36:2), PC(P-33:1), PC(O-33:2) | 0.992424 | 2.20E-09 | 2.2001 |
|  | 1-hexadecanyl-2-(8-[3]-ladderane-octanyl)-sn-glycerophosphocholine | 0.992424 | 1.41E-07 | 1.5856 |
|  | CerP(d44:2) | 0.992424 | 1.48E-06 | 2.1049 |
|  | PE(P-38:0), PE(O-38:1), PC(P-35:0), PC(O-35:1) | 0.992424 | 9.83E-07 | 2.6016 |
|  | PE(38:6), PC(35:6) | 0.992424 | 2.42E-12 | -4.2656 |
|  | SM(d41:2) | 0.992424 | 5.38E-08 | 1.5305 |
|  | SM(d41:1) | 0.992424 | 7.17E-08 | 1.6336 |
|  | PE(42:8), PC(39:8) | 0.992424 | 1.49E-10 | -5.6085 |
|  | PE(42:4), PC(39:4) | 0.992424 | 1.55E-09 | -3.2755 |
|  | PI(36:4) | 0.992424 | 2.10E-07 | -2.3926 |
|  | FA(22:5), DPA | 0.988636 | 2.94E-08 | -2.2968 |
|  | CerP(d34:1) | 0.988636 | 1.60E-08 | 1.2709 |
|  | CerP(d36:1) | 0.988636 | 0.000102 | 1.0385 |
|  | PA(32:0) | 0.988636 | 8.27E-10 | -2.1916 |
|  | PA(33:0) | 0.988636 | 4.03E-11 | -6.3342 |
|  | PA(34:1) | 0.988636 | 3.98E-09 | -2.421 |
|  | PA(35:2) | 0.988636 | 4.00E-09 | -3.044 |
|  | PC(dO-34:4) | 0.988636 | 3.11E-06 | 2.0758 |
|  | PE(40:7), PC(37:7) | 0.988636 | 2.97E-11 | -6.356 |
|  | PI(37:4) | 0.988636 | 1.37E-08 | -2.9727 |
|  | PE(O-18:0), PC(O-15:0) | 0.984848 | 9.57E-07 | 1.6323 |
|  | PA(36:4) | 0.984848 | 4.32E-10 | -1.8955 |
|  | PE(34:0), PC(31:0) | 0.984848 | 2.62E-07 | -1.6921 |
|  | PE(39:4), PC(36:4) | 0.984848 | 6.63E-10 | -3.5431 |
|  | PI(40:5) | 0.984848 | 7.99E-09 | -2.1783 |
|  | PE(38:4), PC(35:4) | 0.981061 | 2.20E-09 | -1.5225 |
|  | PA(34:0) | 0.977273 | 1.41E-07 | -2.8809 |
|  | PS(40:6) | 0.977273 | 3.55E-09 | -4.0082 |
|  | PA(35:1) | 0.973485 | 3.24E-09 | -5.2614 |
|  | PE(34:1), PC(31:1) | 0.973485 | 1.68E-08 | -2.2484 |
|  | PI(36:2) | 0.973485 | 9.74E-08 | 1.6898 |
|  | PI(36:1) | 0.973485 | 1.34E-06 | 1.9049 |
|  | PA(16:0), LPA(16:0) | 0.969697 | 2.08E-07 | -2.1731 |
|  | PA(18:1) | 0.969697 | 3.65E-09 | -2.5084 |
|  | PE(32:0), PC(29:0) | 0.969697 | 1.54E-07 | -3.4461 |
|  | PE(P-36:3), PE(O-36:4) | 0.969697 | 1.50E-07 | 1.2375 |
|  | PE(P-36:2), PE(O-36:3), PC(P-33:2) | 0.969697 | 4.22E-09 | 2.3265 |
|  | PA(40:4) | 0.969697 | 6.75E-08 | -1.8302 |
|  | PA(36:3) | 0.965909 | 1.06E-06 | -1.2249 |
|  | PE(36:1), PC(33:1) | 0.965909 | 1.10E-07 | -2.2765 |
|  | PI(38:4) | 0.965909 | 2.86E-07 | -1.1125 |
|  | PE(38:3), PC(35:3) | 0.962121 | 3.50E-08 | -1.5517 |
|  | PI(39:4) | 0.962121 | 2.99E-08 | -3.1476 |
|  | DHAP(18:0) | 0.958333 | 0.00013 | 1.1481 |
|  | PE(P-34:2), PE(O-34:3) | 0.958333 | 9.25E-08 | 2.7185 |
|  | PG(34:1) | 0.958333 | 2.57E-06 | -1.8094 |
|  | PE(P-38:4), PE(O-38:5) | 0.954545 | 1.30E-05 | 0.70245 |
|  | PE(40:2), PC(37:2) | 0.954545 | 1.31E-07 | -6.4452 |
|  | CPA(18:1) | 0.950758 | 2.36E-07 | -2.7233 |
|  | PE(20:1), LPE(20:1), PC(17:1) | 0.950758 | 2.29E-07 | -4.6813 |
|  | PA(P-38:6) | 0.950758 | 8.78E-08 | -6.7387 |
|  | PE(38:2), PC(35:2) | 0.950758 | 3.12E-06 | -1.5435 |
|  | SM(d39:1) | 0.950758 | 7.76E-05 | 0.73373 |
|  | PE(38:1), PC(35:1) | 0.950758 | 1.07E-06 | -2.3454 |
|  | PE(40:4), PC(37:4) | 0.950758 | 1.76E-08 | -1.908 |
|  | PE(22:4), LPE(22:4) | 0.943182 | 2.86E-07 | -3.1436 |
|  | PA(O-38:4), PA(P-38:3) | 0.943182 | 2.59E-05 | 1.5648 |
|  | PA(38:2) | 0.939394 | 7.70E-07 | -2.8475 |
|  | PG(36:2) | 0.939394 | 3.95E-07 | -6.5423 |
|  | PS(40:5) | 0.939394 | 4.34E-07 | -3.7523 |
|  | PI(40:6) | 0.939394 | 1.89E-06 | -1.8143 |
|  | LPE(22:6) | 0.935606 | 4.35E-07 | -4.9911 |
|  | CerP(d40:1) | 0.935606 | 0.002197 | 0.79044 |
|  | PE(36:3), PC(33:3) | 0.935606 | 8.49E-05 | -1.7107 |
|  | PE(44:8) | 0.935606 | 1.63E-08 | -7.8337 |
|  | PE(P-40:7) | 0.931818 | 2.91E-06 | -3.134 |
|  | PE(39:6), PC(36:6) | 0.92803 | 6.01E-07 | -3.6744 |
|  | PE(18:0), LPE(18:0), PC(15:0) | 0.924242 | 7.02E-07 | -1.3729 |
|  | PE(P-20:0), PC(P-17:0) | 0.924242 | 0.00029 | 1.5093 |
|  | PI(38:5) | 0.924242 | 7.88E-06 | -2.2982 |
|  | PE(38:4(12OH)) | 0.920455 | 6.76E-05 | -3.7157 |
|  | PA(37:3) | 0.916667 | 1.62E-06 | -9.0773 |
|  | PE(P-38:6) | 0.912879 | 4.52E-06 | -2.0364 |
|  | PI(34:0) | 0.912879 | 2.87E-05 | -6.0413 |
|  | PE-Cer(d36:1(2OH)) | 0.909091 | 2.75E-07 | -10.497 |
|  | SM(d39:2) | 0.905303 | 0.004263 | 0.58442 |
|  | PE(P-38:3), PE(O-38:4), PC(O-35:4) | 0.901515 | 8.95E-06 | 1.1152 |
|  | PS(36:2) | 0.901515 | 3.56E-05 | 1.0655 |
|  | PA(37:2) | 0.897727 | 4.76E-05 | -2.9667 |
|  | PI(34:2) | 0.897727 | 5.26E-05 | 1.129 |
|  | PE(39:7), PC(36:7) | 0.886364 | 4.79E-06 | -6.1852 |
|  | PA(20:1) | 0.882576 | 0.000173 | -3.6796 |
|  | PA(38:3) | 0.878788 | 0.000165 | -1.1263 |
|  | PS(44:8) | 0.878788 | 0.000111 | -3.0601 |
|  | PA(34:2) | 0.871212 | 0.001304 | -0.87365 |
|  | PA(38:4) | 0.871212 | 0.000437 | -0.96248 |
|  | PE(40:8) | 0.871212 | 1.58E-06 | -11.171 |
|  | PS(P-38:4), PS(O-38:5) | 0.871212 | 5.44E-05 | -3.13 |
|  | FA(22:4), Adrenic acid | 0.867424 | 0.000175 | -1.1585 |
|  | PE(P-36:4), PE(O-36:5) | 0.867424 | 0.000316 | 0.2604 |
|  | PI(36:3) | 0.863636 | 0.002389 | 0.50568 |
|  | CPA(18:2) | 0.859848 | 0.000467 | -1.1195 |
|  | PE(P-34:1), PE(O-34:2), PC(P-31:1) | 0.859848 | 0.000137 | 0.90188 |
|  | FA(20:1) | 0.856061 | 0.000674 | -5.2029 |
|  | PE(P-40:3), PE(O-40:4), PC(O-37:4) | 0.852273 | 5.96E-05 | 1.0187 |
|  | FA(18:2) | 0.848485 | 0.000304 | 0.43669 |
|  | PA(O-36:4), PA(P-36:3) | 0.848485 | 0.000157 | 1.3334 |
|  | PI(16:0) | 0.840909 | 0.000484 | -3.0943 |
|  | PG(44:11) | 0.837121 | 0.000994 | -2.5706 |
|  | PS(44:6) | 0.837121 | 0.000469 | -3.09 |
|  | PG(36:1) | 0.833333 | 0.000837 | -1.4466 |
|  | PE(18:1), LPE(18:1), PC(15:1) | 0.829545 | 0.001622 | -2.7897 |
|  | PI(40:4) | 0.829545 | 0.00082 | -0.90853 |
|  | PE(44:10) | 0.825758 | 0.000616 | -9.9661 |
|  | PI(O-36:4), PI(P-36:3) | 0.82197 | 0.000473 | 1.0717 |
|  | PI(O-38:4), PI(P-38:3) | 0.818182 | 0.000264 | 0.94008 |
|  | PI(20:4) | 0.818182 | 0.001227 | -1.3123 |
|  | FA(24:4) | 0.814394 | 0.001521 | -3.2509 |
|  | PA(36:1) | 0.810606 | 0.003222 | -1.1106 |
|  | PA(O-38:5), PA(P-38:4) | 0.810606 | 0.000388 | 0.84159 |
|  | PS(40:4) | 0.806818 | 0.002832 | -2.519 |
|  | Cholesterol sulfate | 0.80303 | 0.003932 | 0.97733 |
|  | PA(O-40:5), PA(P-40:4) | 0.80303 | 0.00104 | 0.98933 |
|  | PE(P-40:6) | 0.80303 | 0.002234 | -1.2432 |
|  | PG(44:12) | 0.80303 | 0.001483 | -1.3565 |
|  | PI(40:7) | 0.799242 | 0.011015 | -4.7089 |
|  | SM(d33:2), PE-Cer(d36:2) | 0.799242 | 0.01006 | 0.92725 |
|  | FA(16:0), Palmitic acid | 0.791667 | 0.00298 | -1.048 |
|  | PE(P-16:0) | 0.791667 | 0.003986 | -0.11706 |
|  | PS(40:1) | 0.787879 | 0.008587 | 1.1117 |
|  | PA(20:4) | 0.784091 | 0.001353 | -1.6806 |
|  | PE(20:0), LPE(20:0), PC(17:0), PC(O-17:0) | 0.784091 | 0.013185 | -1.1969 |
|  | PE(O-42:6) | 0.784091 | 0.001716 | 0.7407 |
|  | FA(18:1), Oleic acid | 0.776515 | 0.001807 | -1.6855 |
|  | PE(36:0), PC(33:0) | 0.772727 | 0.005872 | -1.0666 |
|  | PA(O-40:6), PA(P-40:5) | 0.768939 | 0.008637 | 0.50011 |
|  | PE(P-42:4) | 0.765152 | 0.002606 | 0.72817 |
|  | PI(38:2) | 0.765152 | 0.01703 | 0.84919 |
|  | PE(37:2), PC(34:2) | 0.761364 | 0.004981 | -1.4957 |
|  | PS(42:4) | 0.761364 | 0.008584 | 0.65187 |
|  | PE(P-40:4), PE(O-40:5) | 0.753788 | 0.007388 | 0.32034 |
|  | PI(O-38:5), PI(P-38:4) | 0.753788 | 0.007724 | 0.40079 |
|  | PA(36:2) | 0.75 | 0.019426 | -0.86972 |
|  | PE(37:5), PC(34:5) | 0.75 | 0.003444 | -1.1365 |
|  | FA(20:2) | 0.746212 | 0.031371 | -0.96724 |
|  | PE(P-40:5), PE(O-40:6) | 0.742424 | 0.076159 | -0.62929 |
|  | PC(36:2) | 0.734848 | 0.046337 | -4.0561 |
|  | PG(40:7) | 0.731061 | 0.022018 | -0.87242 |
|  | FA(20:3) | 0.723485 | 0.051442 | -0.90393 |
|  | PS(34:1) | 0.723485 | 0.147331 | -1.521 |
|  | CPA(18:0) | 0.719697 | 0.005385 | -0.89183 |
|  | PI(38:6) | 0.719697 | 0.073152 | -1.3473 |
|  | PE(20:2), LPE(20:2), PC(17:2) | 0.715909 | 0.017783 | -1.5886 |
|  | PG(36:0) | 0.712121 | 0.199897 | -0.5896 |
|  | PS(18:0) | 0.708333 | 0.026999 | 1.3913 |
|  | PA(O-36:5), PA(P-36:4) | 0.708333 | 0.080775 | -0.08266 |
|  | PI(40:3) | 0.700758 | 0.155991 | -3.7234 |
|  | PS(38:4) | 0.69697 | 0.063405 | 0.098816 |
|  | PG(44:10) | 0.69697 | 0.033487 | -1.9817 |
|  | PGP(37:0) | 0.689394 | 0.098495 | -2.9055 |
|  | PI(34:1) | 0.685606 | 0.140008 | 0.12511 |
|  | 22:0-Glc-Sitosterol | 0.685606 | 0.116164 | -1.0715 |
|  | PIP(38:4) | 0.685606 | 0.117781 | -1.6133 |
|  | PA(18:0), LPA(18:0) | 0.681818 | 0.056088 | -0.60149 |
|  | PE(39:5), PC(36:5) | 0.681818 | 0.033896 | -0.74981 |
|  | PS(38:1) | 0.67803 | 0.057591 | -4.759 |
|  | Cer(d34:1) | 0.666667 | 0.137271 | 0.46296 |
|  | PS(42:1) | 0.666667 | 0.128899 | 0.004574 |
|  | PG(34:0) | 0.662879 | 0.180296 | -0.72128 |
|  | FA(18:0), Stearic acid | 0.659091 | 0.144082 | -0.68133 |
|  | SM(d37:1), PE-Cer(d40:1) | 0.655303 | 0.39814 | -3.9073 |
|  | PE(P-38:5), PE(O-38:6) | 0.655303 | 0.179099 | -0.72543 |
|  | Cer(d42:2) | 0.651515 | 0.218638 | 0.46683 |
|  | PE(36:2), PC(33:2) | 0.647727 | 0.114005 | -0.5748 |
|  | PS(36:1) | 0.647727 | 0.124035 | -0.78511 |
|  | PS(44:4) | 0.643939 | 0.156574 | 0.34149 |
|  | PS(38:3) | 0.636364 | 0.537416 | 0.11909 |
|  | PG(40:5) | 0.625 | 0.128652 | -1.0993 |
|  | 1-Oleoylglycerophosphoinositol | 0.613636 | 0.86446 | -0.96105 |
|  | FA(20:4), Arachidonic acid | 0.598485 | 0.247581 | -0.51275 |
|  | PS(44:3) | 0.590909 | 0.456029 | 0.54649 |
|  | PG(38:6) | 0.587121 | 0.416805 | 0.09998 |
|  | PG(40:6) | 0.583333 | 0.329974 | -0.64507 |
|  | PE(20:4), LPE(20:4) | 0.579545 | 0.745851 | -0.32368 |
|  | PI(O-40:4), PI(P-40:3) | 0.579545 | 0.975775 | -1.0072 |
|  | PE(P-42:6) | 0.57197 | 0.644296 | -0.69278 |
|  | PG(36:4) | 0.568182 | 0.497601 | 0.32955 |
|  | PG(40:8) | 0.568182 | 0.629704 | -0.21064 |
|  | PG(36:3) | 0.568182 | 0.510763 | -0.1355 |
|  | PG(38:5) | 0.564394 | 0.374477 | -0.35024 |
|  | PG(44:8) | 0.564394 | 0.565878 | -2.6924 |
|  | PC(20:4) | 0.560606 | 0.876093 | -1.9382 |
|  | PI(38:3) | 0.55303 | 0.823066 | -0.73014 |
|  | PA(P-16:0) | 0.541667 | 0.902236 | -0.51422 |
|  | PG(42:10) | 0.541667 | 0.756587 | -0.34735 |
|  | PG(42:9) | 0.537879 | 0.617782 | -0.79914 |
|  | PE(34:2), PC(31:2) | 0.522727 | 0.372429 | -0.58939 |
|  | PS(44:5) | 0.522727 | 0.88885 | -0.3673 |
|  | PI(18:0) | 0.518939 | 0.999766 | -0.50083 |
|  | PG(38:4) | 0.518939 | 0.751671 | -0.08679 |
|  | PG(42:8) | 0.518939 | 0.767588 | -0.88172 |
|  | SM(d35:1), PE-Cer(d38:1) | 0.515152 | 0.441012 | -0.73224 |
|  | PA(O-38:6), PA(P-38:5) | 0.503788 | 0.889487 | -0.38026 |
|  | FA(20:1) | 1 | 9.86E-11 | -5.5478 |
|  | FA(22:5), DPA | 1 | 8.05E-09 | -2.9434 |
|  | FA(24:4) | 1 | 9.22E-10 | -6.1141 |
|  | CPA(16:0) | 1 | 4.70E-16 | -2.4018 |
|  | PA(16:0), LPA(16:0) | 1 | 5.67E-07 | -2.2644 |
|  | CPA(18:2) | 1 | 6.05E-08 | -1.5969 |
|  | CPA(18:1) | 1 | 4.09E-16 | -3.013 |
|  | DHAP(18:0) | 1 | 3.18E-08 | 1.7389 |
|  | PA(18:1) | 1 | 4.62E-11 | -2.6674 |
|  | PE(16:0), LPE(16:0), PC(13:0) | 1 | 7.73E-12 | -4.7586 |
|  | PE(P-18:0), PE(O-18:1), PC(P-15:0) | 1 | 1.94E-13 | 0.62562 |
|  | PE(O-18:0), PC(O-15:0) | 1 | 4.17E-13 | 4.3453 |
|  | PE(18:1), LPE(18:1), PC(15:1) | 1 | 4.10E-06 | -2.9908 |
|  | PE(P-20:0), PC(P-17:0) | 1 | 1.03E-13 | 5.1688 |
|  | PE(20:1), LPE(20:1), PC(17:1) | 1 | 5.63E-10 | -4.4784 |
|  | LPE(22:6) | 1 | 7.69E-10 | -5.1287 |
|  | PE(22:4), LPE(22:4) | 1 | 3.36E-09 | -4.0108 |
|  | CerP(d34:1) | 1 | 2.91E-18 | 1.8944 |
|  | CerP(d34:0) | 1 | 6.07E-12 | 5.0091 |
|  | CerP(d36:1) | 1 | 5.57E-10 | 2.4726 |
|  | PA(32:1) | 1 | 1.85E-14 | -8.6263 |
|  | PA(O-34:2), PA(P-34:1) | 1 | 7.12E-14 | 6.4047 |
|  | PA(O-34:1), PA(P-34:0) | 1 | 4.68E-14 | 4.5049 |
|  | PA(33:0) | 1 | 1.24E-11 | -7.2332 |
|  | PA(34:1) | 1 | 4.85E-14 | -2.4176 |
|  | PA(34:0) | 1 | 3.47E-09 | -3.5442 |
|  | PA(O-36:4), PA(P-36:3) | 1 | 9.41E-14 | 5.0141 |
|  | PA(O-36:3), PA(P-36:2) | 1 | 1.31E-10 | 6.7075 |
|  | PA(35:2) | 1 | 3.25E-10 | -3.5818 |
|  | PA(O-36:2), PA(P-36:1) | 1 | 7.97E-14 | 5.2735 |
|  | PA(35:1) | 1 | 1.16E-11 | -5.5214 |
|  | SM(d33:1), PE-Cer(d36:1) | 1 | 3.50E-16 | 2.2902 |
|  | SM(d33:0) | 1 | 2.65E-13 | 3.6894 |
|  | PE(32:0), PC(29:0) | 1 | 1.40E-09 | -4.0771 |
|  | PA(36:3) | 1 | 1.13E-08 | -1.5151 |
|  | PE(P-34:2), PE(O-34:3) | 1 | 4.54E-12 | 5.0945 |
|  | CerP(d40:1) | 1 | 1.28E-12 | 4.4323 |
|  | PE(P-34:0), PE(O-34:1), PC(P-31:0), PC(O-31:1) | 1 | 6.22E-14 | 2.4587 |
|  | PE-Cer(d36:1(2OH)) | 1 | 9.43E-13 | -9.2968 |
|  | PE(O-34:0), PC(O-31:0) | 1 | 5.01E-14 | 5.7712 |
|  | PA(O-38:5), PA(P-38:4) | 1 | 2.24E-11 | 1.5798 |
|  | PA(37:4) | 1 | 1.57E-12 | -6.2058 |
|  | PA(O-38:4), PA(P-38:3) | 1 | 3.77E-13 | 4.8496 |
|  | PA(37:3) | 1 | 2.35E-15 | -9.8363 |
|  | PA(37:2) | 1 | 4.32E-08 | -3.1018 |
|  | PE(34:1), PC(31:1) | 1 | 3.15E-13 | -2.3746 |
|  | PA(38:6) | 1 | 5.07E-12 | -3.8801 |
|  | PA(38:5) | 1 | 1.54E-12 | -2.855 |
|  | PE(P-36:3), PE(O-36:4) | 1 | 4.78E-12 | 2.1242 |
|  | PC(dO-34:4) | 1 | 7.68E-14 | 4.9326 |
|  | PE(P-36:2), PE(O-36:3), PC(P-33:2) | 1 | 3.84E-13 | 4.1235 |
|  | CerP(d42:2) | 1 | 2.56E-16 | 2.615 |
|  | PA(38:2) | 1 | 8.03E-09 | -2.736 |
|  | PE(P-36:1), PE(O-36:2), PC(P-33:1), PC(O-33:2) | 1 | 1.98E-15 | 4.4656 |
|  | CerP(d42:1) | 1 | 2.62E-13 | 4.1985 |
|  | PE(P-36:0), PE(O-36:1), PC(P-33:0), PC(O-33:1) | 1 | 1.53E-17 | 3.5224 |
|  | PA(O-40:5), PA(P-40:4) | 1 | 4.09E-10 | 3.1263 |
|  | PA(39:4) | 1 | 1.05E-11 | -4.3674 |
|  | PE(36:4), PE(P-36:4), PC(33:4) | 1 | 1.08E-16 | -2.5025 |
|  | PE(36:3), PC(33:3) | 1 | 1.40E-05 | -2.4379 |
|  | PE(36:1), PC(33:1) | 1 | 3.13E-13 | -2.0422 |
|  | PA(40:7) | 1 | 9.18E-16 | -6.2344 |
|  | PA(40:6) | 1 | 7.90E-12 | -3.0645 |
|  | PA(40:5) | 1 | 4.94E-11 | -2.4346 |
|  | PA(40:4) | 1 | 3.28E-09 | -2.251 |
|  | PE(P-38:3), PE(O-38:4), PC(O-35:4) | 1 | 1.60E-15 | 2.4932 |
|  | 1-hexadecanyl-2-(8-[3]-ladderane-octanyl)-sn-glycerophosphocholine | 1 | 7.31E-11 | 2.7449 |
|  | PE(P-38:2) | 1 | 9.32E-11 | 4.7075 |
|  | CerP(d44:2) | 1 | 2.29E-10 | 3.4897 |
|  | PE(P-38:1), PE(O-38:2), PC(P-35:1), PC(O-35:2) | 1 | 6.31E-10 | 5.5112 |
|  | PE(P-38:0), PE(O-38:1), PC(P-35:0), PC(O-35:1) | 1 | 1.25E-10 | 8.018 |
|  | PE(38:6), PC(35:6) | 1 | 5.64E-12 | -4.173 |
|  | PE(38:5), PC(35:5) | 1 | 9.02E-18 | -3.4274 |
|  | PE(38:4), PC(35:4) | 1 | 1.83E-14 | -1.7402 |
|  | PE(38:3), PC(35:3) | 1 | 3.36E-07 | -1.7783 |
|  | SM(d39:1) | 1 | 8.19E-11 | 2.4033 |
|  | PE(P-40:7) | 1 | 5.30E-11 | -2.4091 |
|  | PA(42:7) | 1 | 1.42E-12 | -8.2313 |
|  | PG(36:2) | 1 | 1.49E-09 | -5.5118 |
|  | PE(P-40:3), PE(O-40:4), PC(O-37:4) | 1 | 1.97E-11 | 2.1895 |
|  | PE(40:8) | 1 | 3.67E-16 | -12.534 |
|  | PE(40:7), PC(37:7) | 1 | 9.58E-12 | -6.6043 |
|  | PE(40:6), PC(37:6) | 1 | 4.96E-18 | -2.7457 |
|  | PE(40:5), PC(37:5) | 1 | 3.35E-17 | -2.8295 |
|  | PE(40:3), PC(37:3) | 1 | 1.65E-12 | -3.1715 |
|  | SM(d41:2) | 1 | 4.49E-15 | 2.7916 |
|  | PE(40:2), PC(37:2) | 1 | 5.20E-13 | -5.9202 |
|  | SM(d41:1) | 1 | 3.09E-13 | 2.8337 |
|  | PE(O-42:6) | 1 | 2.94E-12 | 2.8675 |
|  | PE(P-42:4) | 1 | 3.70E-11 | 3.458 |
|  | PE(42:8), PC(39:8) | 1 | 1.18E-11 | -5.8964 |
|  | PE(42:7), PC(39:7) | 1 | 1.44E-14 | -7.5401 |
|  | PE(42:6), PC(39:6) | 1 | 3.54E-12 | -3.4804 |
|  | PE(42:5), PC(39:5) | 1 | 1.25E-14 | -5.3574 |
|  | PE(42:4), PC(39:4) | 1 | 1.54E-09 | -3.3777 |
|  | PS(40:6) | 1 | 9.51E-10 | -4.4563 |
|  | PS(40:5) | 1 | 1.08E-09 | -4.9684 |
|  | PI(34:0) | 1 | 5.08E-11 | -7.8505 |
|  | PE(44:10) | 1 | 5.58E-13 | -8.524 |
|  | PS(40:4) | 1 | 5.63E-09 | -3.4349 |
|  | PE(44:8) | 1 | 4.29E-16 | -8.9167 |
|  | PI(O-36:4), PI(P-36:3) | 1 | 5.48E-10 | 4.1547 |
|  | PI(36:4) | 1 | 3.97E-13 | -2.6227 |
|  | PI(36:2) | 1 | 8.69E-18 | 2.6267 |
|  | PI(36:1) | 1 | 4.50E-17 | 3.2664 |
|  | PI(O-38:5), PI(P-38:4) | 1 | 1.86E-08 | 1.7815 |
|  | PI(O-38:4), PI(P-38:3) | 1 | 1.08E-08 | 2.6488 |
|  | PI(38:5) | 1 | 7.09E-15 | -2.5929 |
|  | PI(38:4) | 1 | 1.34E-11 | -1.3959 |
|  | PI(40:6) | 1 | 5.14E-10 | -2.7361 |
|  | PI(40:5) | 1 | 2.65E-12 | -3.0374 |
|  | PI(40:4) | 1 | 1.60E-11 | -1.7291 |
|  | FA(22:6), DHA | 0.994792 | 3.25E-09 | -2.4295 |
|  | PA(36:4) | 0.994792 | 1.07E-10 | -2.0931 |
|  | PS(36:2) | 0.994792 | 3.82E-08 | 1.2236 |
|  | FA(18:1), Oleic acid | 0.989583 | 4.01E-07 | -1.5494 |
|  | Cholesterol sulfate | 0.989583 | 5.16E-06 | 3.0932 |
|  | PA(38:4) | 0.989583 | 6.80E-08 | -1.0711 |
|  | PA(O-40:6), PA(P-40:5) | 0.989583 | 1.34E-06 | 1.3251 |
|  | PE(P-38:4), PE(O-38:5) | 0.989583 | 1.39E-10 | 0.49338 |
|  | SM(d39:2) | 0.989583 | 1.66E-07 | 3.2282 |
|  | PE(39:6), PC(36:6) | 0.989583 | 8.76E-08 | -4.2923 |
|  | PE(39:4), PC(36:4) | 0.989583 | 6.40E-09 | -3.5446 |
|  | PG(44:11) | 0.989583 | 1.56E-07 | -3.1866 |
|  | PI(40:3) | 0.989583 | 5.17E-08 | -5.7736 |
|  | FA(22:4), Adrenic acid | 0.984375 | 1.35E-09 | -2.1892 |
|  | PI(16:0) | 0.984375 | 5.49E-07 | -3.5986 |
|  | PA(P-38:6) | 0.984375 | 2.50E-08 | -5.7365 |
|  | PE(P-38:6) | 0.984375 | 2.13E-08 | -1.6078 |
|  | PE(38:1), PC(35:1) | 0.984375 | 1.82E-07 | -1.6623 |
|  | PE(39:7), PC(36:7) | 0.984375 | 1.01E-10 | -6.2338 |
|  | PA(32:0) | 0.979167 | 4.41E-09 | -2.3806 |
|  | PA(38:3) | 0.979167 | 2.20E-06 | -1.1282 |
|  | PG(36:1) | 0.979167 | 2.50E-07 | -1.3128 |
|  | PE(38:4(12OH)) | 0.979167 | 2.35E-08 | -5.6536 |
|  | PGP(37:0) | 0.979167 | 1.27E-07 | -4.1994 |
|  | PI(40:7) | 0.979167 | 2.55E-07 | -5.8072 |
|  | PG(34:1) | 0.973958 | 1.46E-06 | -2.0372 |
|  | PI(37:4) | 0.973958 | 4.47E-08 | -3.3015 |
|  | PE(P-40:6) | 0.96875 | 1.23E-06 | -1.0822 |
|  | PI(39:4) | 0.96875 | 6.85E-08 | -3.3174 |
|  | CPA(18:0) | 0.963542 | 1.72E-05 | -1.1955 |
|  | PE(40:4), PC(37:4) | 0.963542 | 5.34E-10 | -1.8712 |
|  | PS(44:8) | 0.963542 | 1.96E-07 | -3.4097 |
|  | PE(34:0), PC(31:0) | 0.958333 | 1.49E-07 | -1.8019 |
|  | PS(P-38:4), PS(O-38:5) | 0.958333 | 3.81E-07 | -4.1315 |
|  | PE(P-40:4), PE(O-40:5) | 0.953125 | 1.41E-06 | 0.67821 |
|  | PA(34:2) | 0.947917 | 2.65E-05 | -0.94664 |
|  | SM(d35:1), PE-Cer(d38:1) | 0.947917 | 1.58E-05 | 1.9279 |
|  | PS(38:1) | 0.947917 | 1.52E-06 | -6.4065 |
|  | PS(42:4) | 0.947917 | 0.000182 | 1.7042 |
|  | PI(38:6) | 0.947917 | 8.41E-05 | -1.9863 |
|  | PE(18:0), LPE(18:0), PC(15:0) | 0.942708 | 1.68E-07 | -1.3319 |
|  | FA(16:0), Palmitic acid | 0.9375 | 1.43E-06 | -1.2352 |
|  | PI(20:4) | 0.9375 | 1.01E-05 | -1.4315 |
|  | PE(36:0), PC(33:0) | 0.9375 | 2.73E-05 | -1.2967 |
|  | PA(20:1) | 0.932292 | 2.28E-05 | -3.2526 |
|  | PI(34:2) | 0.932292 | 5.11E-06 | 1.5042 |
|  | PG(44:10) | 0.927083 | 9.62E-06 | -2.5769 |
|  | PIP(38:4) | 0.921875 | 1.67E-05 | -4.497 |
|  | FA(20:2) | 0.916667 | 0.000128 | -1.5464 |
|  | PE(P-34:1), PE(O-34:2), PC(P-31:1) | 0.916667 | 7.46E-07 | 1.4352 |
|  | FA(18:2) | 0.911458 | 2.72E-05 | 0.52903 |
|  | PS(34:1) | 0.885417 | 0.000732 | -2.1738 |
|  | FA(20:3) | 0.875 | 0.001245 | -1.2451 |
|  | PE(38:2), PC(35:2) | 0.875 | 0.000182 | -1.1068 |
|  | PI(36:3) | 0.875 | 0.000738 | 0.54006 |
|  | PE(37:5), PC(34:5) | 0.869792 | 0.000108 | -1.9603 |
|  | PS(42:1) | 0.869792 | 0.000322 | -4.0504 |
|  | 1-Oleoylglycerophosphoinositol | 0.864583 | 0.000294 | -1.1828 |
|  | PE(P-42:6) | 0.848958 | 0.000333 | 0.94044 |
|  | PA(20:4) | 0.838542 | 0.000522 | -2.0126 |
|  | PE(P-36:4), PE(O-36:5) | 0.838542 | 0.000374 | 0.041852 |
|  | PG(44:8) | 0.828125 | 0.000145 | -2.0358 |
|  | PE(39:5), PC(36:5) | 0.822917 | 0.001326 | -1.5728 |
|  | PE(20:2), LPE(20:2), PC(17:2) | 0.817708 | 0.003607 | -1.842 |
|  | PA(36:1) | 0.8125 | 0.000391 | -1.0356 |
|  | PA(18:0), LPA(18:0) | 0.802083 | 0.009464 | -0.70849 |
|  | PE(20:4), LPE(20:4) | 0.786458 | 0.005017 | -1.0332 |
|  | PA(36:2) | 0.78125 | 0.002839 | -0.74026 |
|  | PA(O-36:5), PA(P-36:4) | 0.776042 | 0.009407 | -0.04396 |
|  | PI(34:1) | 0.765625 | 0.089747 | 0.4626 |
|  | PI(38:3) | 0.765625 | 0.007796 | -1.1559 |
|  | PS(44:4) | 0.75 | 0.041556 | 0.68787 |
|  | FA(18:0), Stearic acid | 0.744792 | 0.017973 | -0.90022 |
|  | PC(20:4) | 0.744792 | 0.056107 | -2.1803 |
|  | PE(P-38:5), PE(O-38:6) | 0.744792 | 0.017438 | -0.68986 |
|  | PS(44:6) | 0.744792 | 0.097611 | -1.9722 |
|  | PI(18:0) | 0.729167 | 0.05603 | -0.77175 |
|  | PS(18:0) | 0.723958 | 0.019985 | 1.7542 |
|  | PS(44:3) | 0.723958 | 0.102926 | 1.5237 |
|  | PG(34:0) | 0.723958 | 0.022779 | -0.68435 |
|  | PE(P-16:0) | 0.71875 | 0.047005 | -0.21577 |
|  | Cer(d42:2) | 0.71875 | 0.0387 | -1.5989 |
|  | Cer(d34:1) | 0.713542 | 0.048117 | -1.6266 |
|  | PS(36:1) | 0.713542 | 0.020766 | -0.85695 |
|  | SM(d33:2), PE-Cer(d36:2) | 0.708333 | 0.057489 | 0.36701 |
|  | PC(36:2) | 0.697917 | 0.048464 | -2.6394 |
|  | PG(40:5) | 0.697917 | 0.002476 | -1.1066 |
|  | PS(38:4) | 0.692708 | 0.102389 | -0.14345 |
|  | 22:0-Glc-Sitosterol | 0.6875 | 0.098025 | -2.4653 |
|  | PG(42:9) | 0.677083 | 0.005463 | -0.98272 |
|  | PG(40:7) | 0.666667 | 0.02653 | -0.61258 |
|  | PG(40:6) | 0.666667 | 0.020057 | -0.54281 |
|  | PG(42:8) | 0.666667 | 0.03067 | -0.35293 |
|  | PS(44:5) | 0.661458 | 0.302315 | 0.046432 |
|  | PG(38:5) | 0.651042 | 0.252444 | 0.46053 |
|  | PG(38:6) | 0.640625 | 0.18395 | 1.707 |
|  | FA(20:4), Arachidonic acid | 0.635417 | 0.174755 | -0.65497 |
|  | PG(36:0) | 0.630208 | 0.40419 | -0.49079 |
|  | PE(P-40:5), PE(O-40:6) | 0.625 | 0.311533 | -0.51652 |
|  | PI(38:2) | 0.614583 | 0.335071 | -0.45562 |
|  | PE(37:2), PC(34:2) | 0.604167 | 0.3977 | -1.02 |
|  | PS(38:3) | 0.604167 | 0.866789 | -0.16129 |
|  | PA(O-38:6), PA(P-38:5) | 0.598958 | 0.229269 | -0.03552 |
|  | PG(36:3) | 0.583333 | 0.766687 | 0.32015 |
|  | PG(38:4) | 0.578125 | 0.744954 | 0.66948 |
|  | PG(36:4) | 0.5625 | 0.18024 | 1.1354 |
|  | PE(34:2), PC(31:2) | 0.546875 | 0.162607 | -0.88505 |
|  | PG(40:8) | 0.546875 | 0.325172 | 1.0399 |
|  | PS(40:1) | 0.536458 | 0.551839 | -0.82848 |
|  | PE(20:0), LPE(20:0), PC(17:0), PC(O-17:0) | 0.53125 | 0.308039 | -0.41917 |
|  | PI(O-40:4), PI(P-40:3) | 0.526042 | 0.637695 | -0.5387 |
|  | PA(P-16:0) | 0.520833 | 0.937482 | -0.4727 |
|  | PG(42:10) | 0.515625 | 0.555182 | 0.69179 |
|  | PE(36:2), PC(33:2) | 0.505208 | 0.866446 | -0.56493 |
|  | SM(d37:1), PE-Cer(d40:1) | 0.505208 | 0.469099 | -0.98175 |
|  | PG(44:12) | 0.5 | 0.656302 | -0.54388 |

**Cer(d34:1)**, *m/z* 536.5043

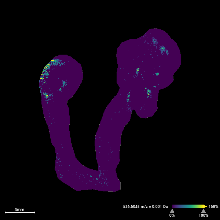

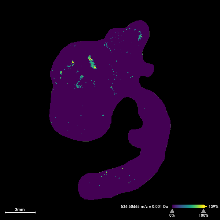

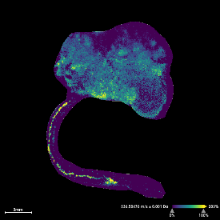

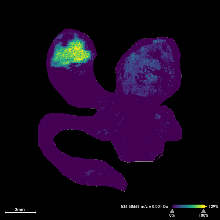

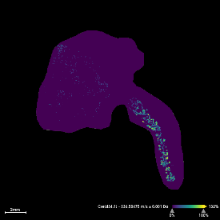

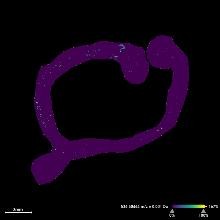

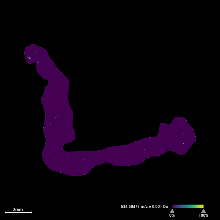

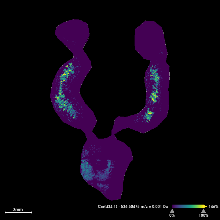

| DKO-2  0.99 ppm | DKO-3  0.71 ppm | TKO-1  0.92 ppm | TKO-2  0.77 ppm | TKO-3  0.90 ppm | Control-1  0.69 ppm | Control-2  0.93 ppm | Control-4  0.90 ppm |
| --- | --- | --- | --- | --- | --- | --- | --- |

**SM(d35:1)**, *m/z* 715.5754

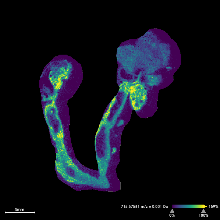

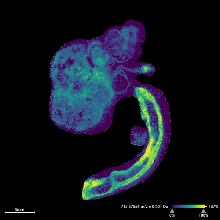

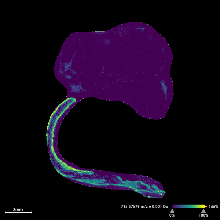

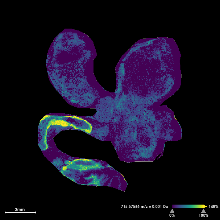

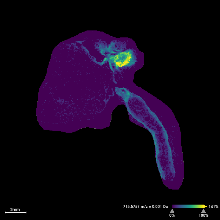

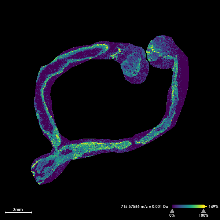

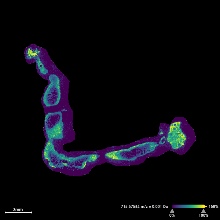

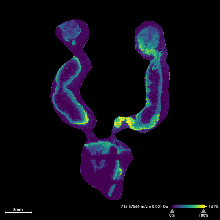

| DKO-2  0.57 ppm | DKO-3  0.22 ppm | TKO-1  0.75 ppm | TKO-2  0.64 ppm | TKO-3  0.42 ppm | Control-1  0.64 ppm | Control-2  0.61 ppm | Control-4  0.36 ppm |
| --- | --- | --- | --- | --- | --- | --- | --- |

**PA(38:5)**, *m/z* 721.4808

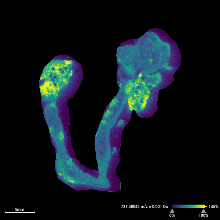

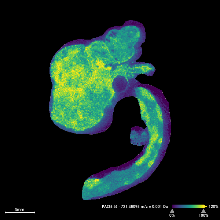

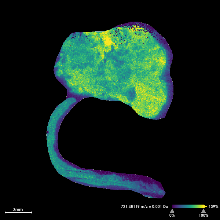

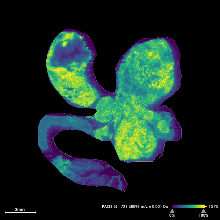

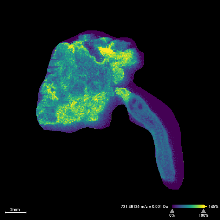

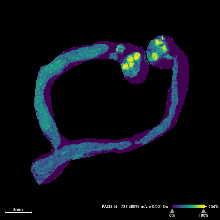

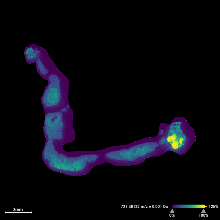

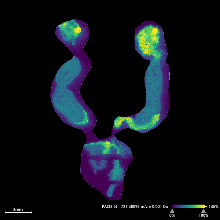

| DKO-2  -0.004 ppm | DKO-3  0.20 ppm | TKO-1  0.49 ppm | TKO-2  0.20 ppm | TKO-3  0.20 ppm | Control-1  0.20 ppm | Control-2  0.54 ppm | Control-4  0.20 ppm |
| --- | --- | --- | --- | --- | --- | --- | --- |

**PA(O-38:4)**, *m/z* 709.5172

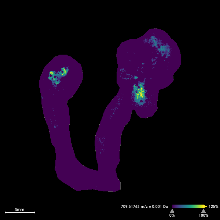

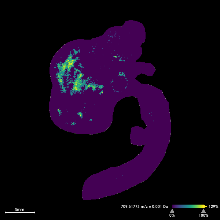

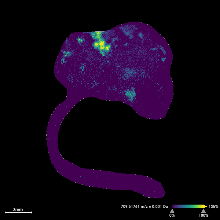

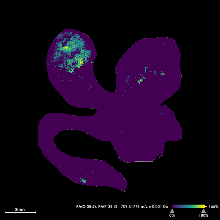

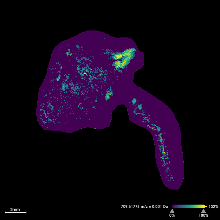

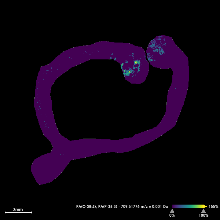

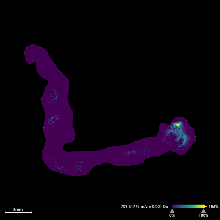

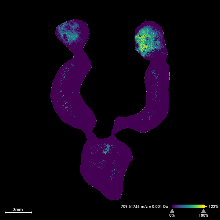

| DKO-2  0.58 ppm | DKO-3  0.72 ppm | TKO-1  0.55 ppm | TKO-2  0.76 ppm | TKO-3  0.79 ppm | Control-1  0.76 ppm | Control-2  0.74 ppm | Control-4  0.37 ppm |
| --- | --- | --- | --- | --- | --- | --- | --- |

**PE(38:1)**, *m/z* 772.5856

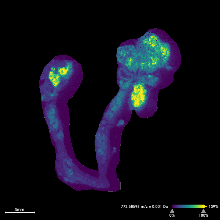

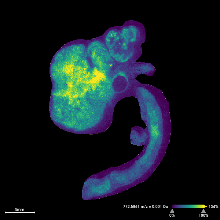

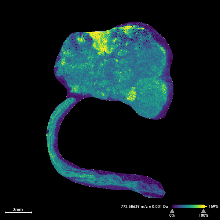

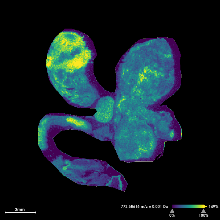

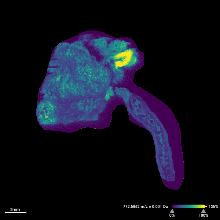

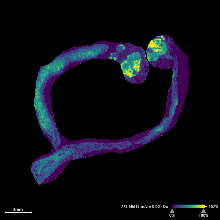

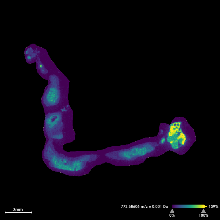

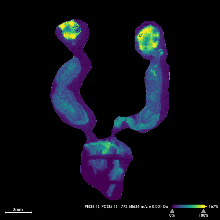

| DKO-2  0.45 ppm | DKO-3  0.94 ppm | TKO-1  0.98 ppm | TKO-2  0.68 ppm | TKO-3  0.74 ppm | Control-1  0.71 ppm | Control-2  0.55 ppm | Control-4  0.94 ppm |
| --- | --- | --- | --- | --- | --- | --- | --- |

**PE(O-34:2)**, *m/z* 700.5281

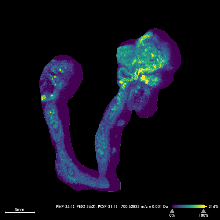

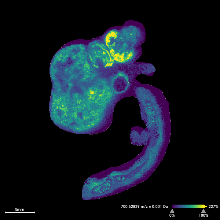

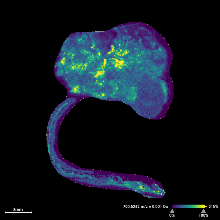

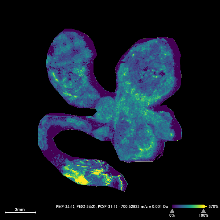

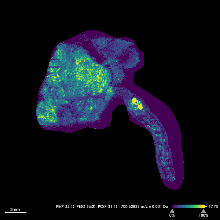

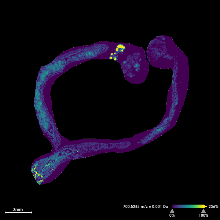

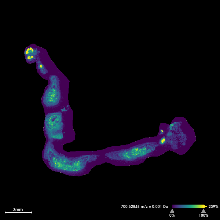

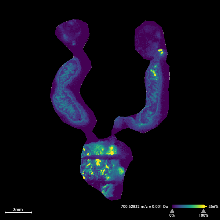

| DKO-2  0.16 ppm | DKO-3  0.25 ppm | TKO-1  0.12 ppm | TKO-2  0.16 ppm | TKO-3  0.16 ppm | Control-1  0.55 ppm | Control-2  0.52 ppm | Control-4  0.29 ppm |
| --- | --- | --- | --- | --- | --- | --- | --- |

**PE(O-40:6)**, *m/z* 776.5594

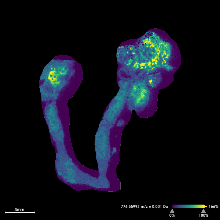

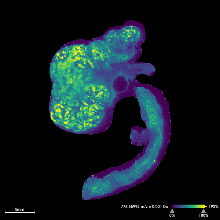

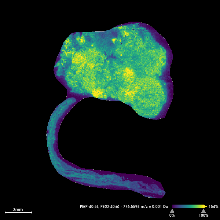

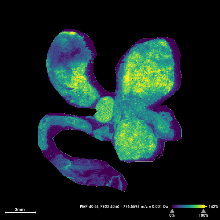

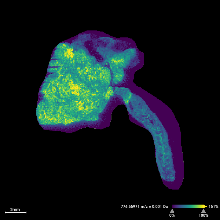

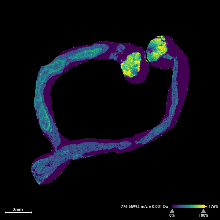

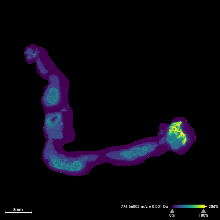

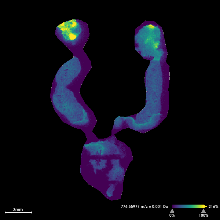

| DKO-2  0.66 ppm | DKO-3  0.67 ppm | TKO-1  0.49 ppm | TKO-2  0.49 ppm | TKO-3  0.38 ppm | Control-1  0.67 ppm | Control-2  0.78 ppm | Control-4  0.46 ppm |
| --- | --- | --- | --- | --- | --- | --- | --- |

**PS(36:2)**, *m/z* 786.5285

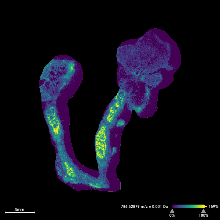

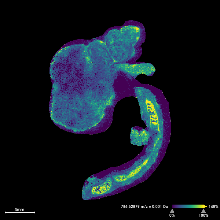

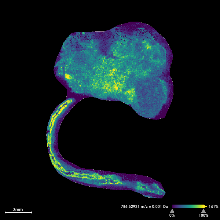

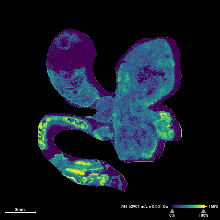

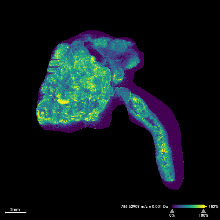

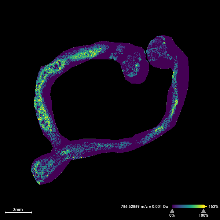

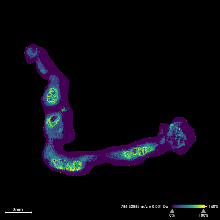

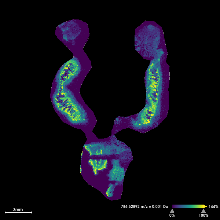

| DKO-2  0.35 ppm | DKO-3  0.35 ppm | TKO-1  0.87 ppm | TKO-2  0.63 ppm | TKO-3  0.73 ppm | Control-1  0.48 ppm | Control-2  0.43 ppm | Control-4  0.52 ppm |
| --- | --- | --- | --- | --- | --- | --- | --- |

**PG(38:5)**, *m/z* 795.5176

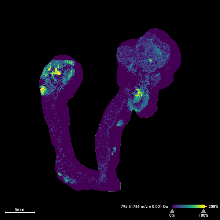

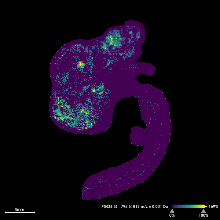

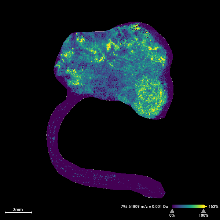

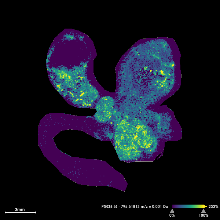

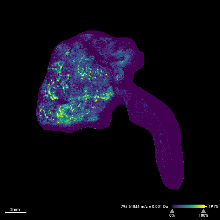

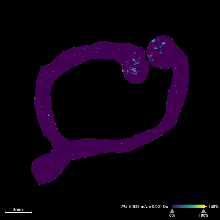

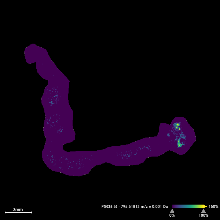

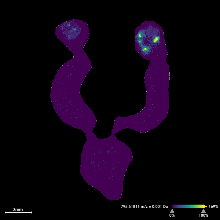

| DKO-2  0.31 ppm | DKO-3  0.65 ppm | TKO-1  0.60 ppm | TKO-2  0.65 ppm | TKO-3  1.06 ppm | Control-1  0.84 ppm | Control-2  0.65 ppm | Control-4  0.62 ppm |
| --- | --- | --- | --- | --- | --- | --- | --- |

**PI(38:4)**, *m/z* 885.5493

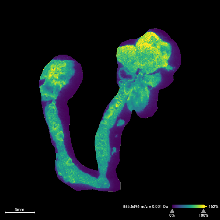

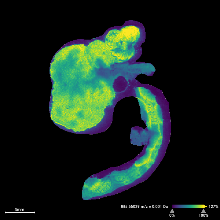

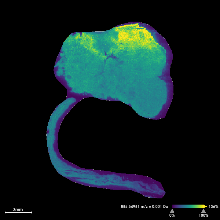

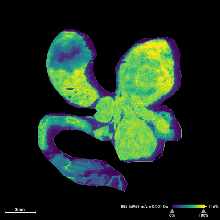

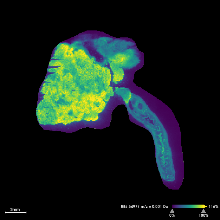

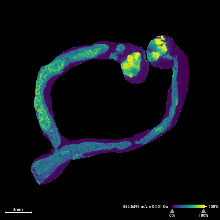

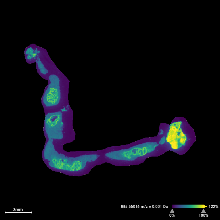

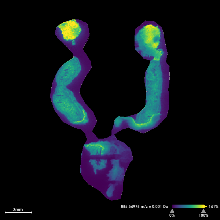

| DKO-2  0.33 ppm | DKO-3  1.11 ppm | TKO-1  0.24 ppm | TKO-2  0.32 ppm | TKO-3  0.24 ppm | Control-1  0.56 ppm | Control-2  0.96 ppm | Control-4  0.53 ppm |
| --- | --- | --- | --- | --- | --- | --- | --- |

**Figure S1**. Selected ion images and their mass errors of the remaining tissue sections studied by MSI that were not displayed in Figure 4 of the main text.

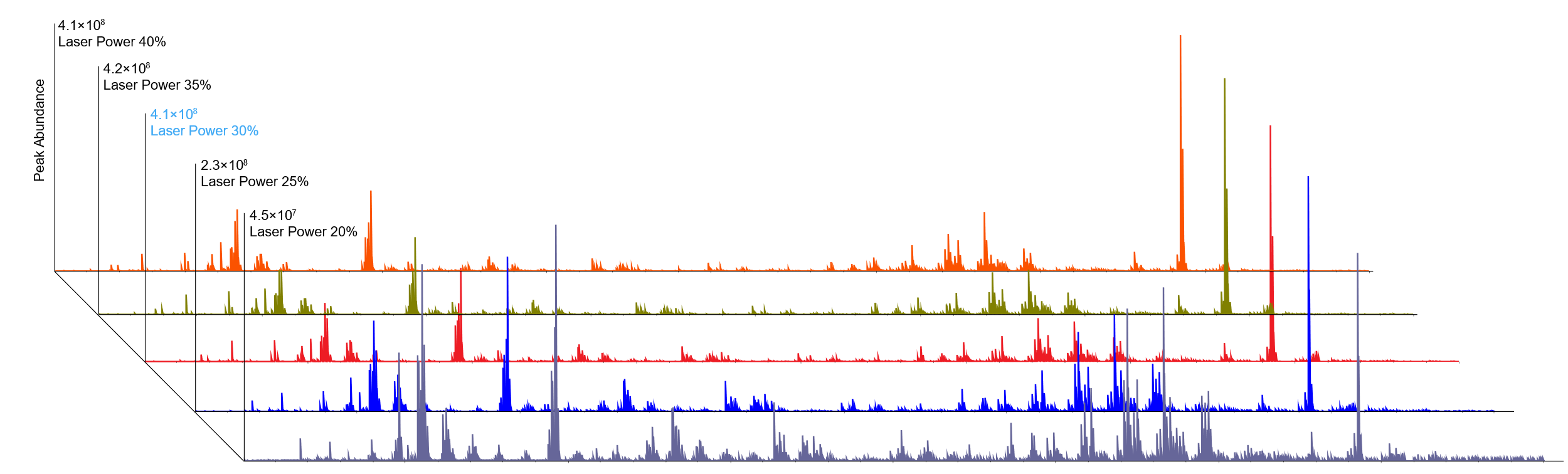

| **Laser Power** | **20%** | **25%** | **30%** | **35%** | **40%** |
| --- | --- | --- | --- | --- | --- |
| PS(42:1) : PA(42:1) | NA | 1.18 | 1.67 | 1.79 | 1.42 |
| PS(40:4) : PA(40:4) | 0.96 | 0.76 | 0.77 | 0.53 | 0.62 |
| PS(40:5) : PA(40:5) | 0.54 | 0.40 | 0.48 | 0.31 | 0.30 |
| PS(40:6) : PA(40:6) | 0.85 | 0.61 | 0.72 | 0.45 | 0.37 |
| PS(38:4) : PA(38:4) | 0.44 | 0.38 | 0.30 | 0.22 | 0.24 |
| PS(P-38:4) : PA(P-38:4) | 0.56 | 0.50 | 0.52 | 0.45 | 0.60 |
| PS(36:1) : PA(36:1) | 1.27 | 1.27 | 0.99 | 1.02 | 1.25 |
| PS(36:2) : PA(36:2) | 0.26 | 0.23 | 0.10 | 0.12 | 0.17 |

**Figure S2**. Mean mass spectra and PS:PA peak area ratios at different laser powers. 30% Laser power (in blue) was used for all final experiments.
